# Human Rab small GTPase- and class V myosin-mediated membrane tethering in a chemically defined reconstitution system

**DOI:** 10.1101/129072

**Authors:** Motoki Inoshita, Joji Mima

**Affiliations:** From the Institute for Protein Reseorch, Osoko University, Suito, Osoko 565-0871, Japan

**Keywords:** Liposome, membrane reconstitution, membrane tethering, membrane trafficking, Myo5, myosin, Rab, small GTPase

## Abstract

Membrane tethering is a fundamental process essential for compartmental specificity of intracellular membrane trafficking in eukaryotic cells. Rab-family small GTPases and specific sets of Rab-interacting effector proteins, including coiled-coil tethering proteins and multisubunit tethering complexes, have been reported to be responsible for membrane tethering. However, whether and how these key components directly and specifically tether subcellular membranes still remains enigmatic. Using chemically defined proteoliposomal systems reconstituted with purified human Rab proteins and synthetic liposomal membranes to study the molecular basis of membrane tethering, we established here that Rab-family GTPases have a highly conserved function to directly mediate membrane tethering, even in the absence of any types of Rab effectors such as the so-called tethering proteins. Moreover, we demonstrate that membrane tethering mediated by endosomal Rab11a is drastically and selectively stimulated by its cognate Rab effectors, class V myosins (Myo5A and Myo5B), in a GTP-dependent manner. Of note, Myo5A and Myo5B exclusively recognized and cooperated with the membrane-anchored form of their cognate Rab11a to support membrane tethering mediated by *trans*-Rab assemblies on apposing membranes. Our findings support the novel concept that Rab-family proteins provide a *bona fide* membrane tether to physically and specifically link two distinct lipid bilayers of subcellular membranes. They further indicate that Rab-interacting effector proteins, including class V myosins, can regulate these Rab-mediated membrane tethering reactions.

## Introduction

All eukaryotic cells, from yeast to human cells, deliver the collect sets of the cargoes such as proteins and lipids to the appropriate cellular compartments including a variety of subcellular organelles, the plasma membrane, and the extracellular space (1). These membrane trafficking events are a fundamental and highly selective process in the eukaryotic endomembrane systems, ensuring that transport vesicles or other membrane-bounded carriers specifically recognize, physically bind to, and eventually fuse with the target membranes in a spatiotemporally regulated manner (1). A large body of prior genetic and biochemical studies have described miscellaneous key protein components functioning in eukaryotic membrane trafficking, which include SNAREs (soluble *N*-ethylmaleimide-sensitive factor attachment protein receptors) (2), SNARE-binding cofactors and chaperones such as Sec1/Munc18 proteins (3), Rab-family small GTPases (4, 5), and Rab-interacting proteins termed “Rab effectors” (6, 7). Membrane tethering mediated by Rab GTPases and Rab effectors is generally known to be the first contact between the transport carriers and the target membranes and thus an essential step to determine the specificity of intracellular membrane trafficking (8, 9, 10), followed by SNARE-mediated membrane fusion, which is another critical layer to confer the fidelity of membrane trafficking (11–15). However, how Rab GTPases and Rab effectors work together to specifically drive membrane tethering has still remained elusive (16–18), although recent biochemical reconstitution studies have begun to report the intrinsic membrane tethering potency of specific Rab effectors (10, 19–23) and Rab GTPases (16, 18). In this study, to gain a deeper insight into the mechanisms of intracellular membrane tethering, we have reconstituted membrane tethering reactions in a chemically defined system from synthetic liposomes and purified human Rab GTPases and class V myosins as the cognate effectors of Rab11a, thereby comprehensively investigating their genuine functions in membrane tethering.

## Results and discussion

Prior two studies on membrane tethering in a chemically defined reconstitution system have described that tethering of synthetic liposomes can be directly triggered by several Rab-family GTPases themselves, which locate and function at the endosomal compartments in yeast (16) and human cells (18). This Rab-mediated tethering of liposomal membranes is an efficient and specific biochemical reaction, as we established that membrane-anchored human Rab5a rapidly induced the formation of massive liposome clusters, and also that the tethering activity of Rab5a was able to be strictly and reversibly controlled by the membrane attachment and detachment of Rab proteins on both apposing membranes (18). Intriguingly, Rab5a-mediated membrane tethering was not blocked by addition of a large excess of soluble Rab5a proteins that lack a membrane anchor at their C-terminus (18), indicating that this membrane tethering process cannot be explained simply by protein dimerization and protein-protein interactions in solution. These our results strongly suggest the critical requirement of *trans*-Rab-Rab interactions on two distinct opposing membranes for a reversible membrane tethering event (18).

In the current reconstitution study, to further comprehensively investigate the inherent membrane tethering potency of Rab-family GTPases in human, we purified and tested the six representative Rab GTPases functioning in the endocytic pathways (Rab4a, Rab5a, Rab7a, Rab9a, Rab11a, and Rab14) (7), for typical non-endosomal Rabs, Rab1a in ER-to-Golgi traffic and Rab3a in exocytosis (4, 5), and also HRas for the control protein of a non-Rab small GTPase in the Ras superfamily (24) (Fig. 1A). All of the Rab-family and HRas GTPase proteins were purified as their full-length forms, consisting of the N-terminal nonconserved flexible segments (5-30 residues), the conserved globular Ras-superfamily GTPase domains in the middle (160-170 residues), and the so-called C-terminal hypervariable region (HVR) domains (20-50 residues) (24) (Fig. 1A,B). In addition to the full-length amino acid sequences of native proteins, to mimic membrane binding and anchoring of native Rab and HRas proteins through their isoprenyl or palmitoyl lipid anchors at the C-terminus (5, 24), these recombinant Rab and HRas proteins used here were further artificially modified with a C-terminal polyhistidine tag (His12), which can be stably associated with synthetic liposomal membranes bearing a DOGS-NTA lipid (1,2-dioleoyl-sn-glycero-3-{[N-(5-amino-1-carboxypentyl)iminodiacetic acid]-succinyl}) (Fig. 1B; Fig. 2A). We employed GTPase activity assays for all of the purified Rab-His12 and HRas-His12 proteins, indicating that they retained the comparable intrinsic GTPase activities, which specifically converted GTP to GDP and a free phosphate (Fig. 1C). Thus, this establishes that purified Rab-His12 and HRas-His12 proteins from the current preparations are all well folded and functionally active in solution (Fig. 1C).

**Figure 1.**
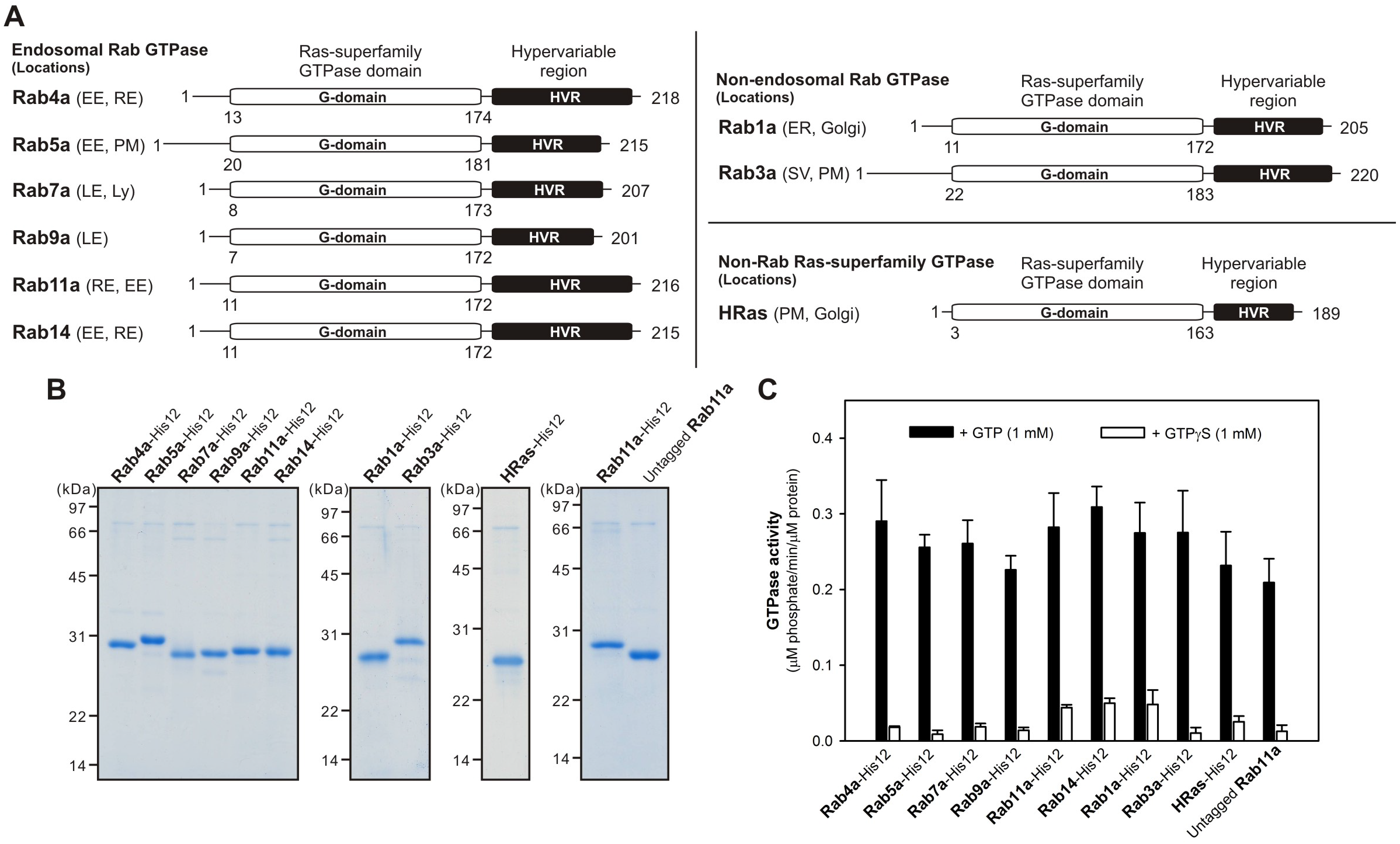
Purification of human Rab GTPases used in this reconstitution study. (**A**) Schematic representation of the endosomal Rabs (Rab4a, Rab5a, Rab7a, Rab9a, Rab11a, and Rab14), non-endosomal Rabs (Rab1a and Rab3a), and HRas GTPase in human, showing their amino acid residues, domains (Ras-superfamily GTPase domains and hypervariable regions), and intracellular locations, which include early endosome (EE), recycling endosome (RE), plasma membrane (PM), late endosome (LE), lysosome (Ly), endoplasmic reticulum (ER), Golgi, and secretory vesicle (SV). (**B**) Coomassie Blue-stained gels of purified recombinant proteins of the C-terminally His12-tagged endosomal Rab, non-endosomal Rab, and HRas GTPases and the untagged form of Rab11a used in this study. (**C**) Intrinsic GTP-hydrolysis activities of purified Rab proteins. Purified Rab-His12, HRas-His12, and untagged Rab11a proteins (2 μM) were incubated at 30°C for 1 h in RB150 containing MgCl_2_ (6 mM), DTT (1 mM), and GTP (1 mM) or GTPγS (1 mM) for the control, followed by assaying released free phosphate molecules using a Malachite Green-based reagent.

**Figure 2.**
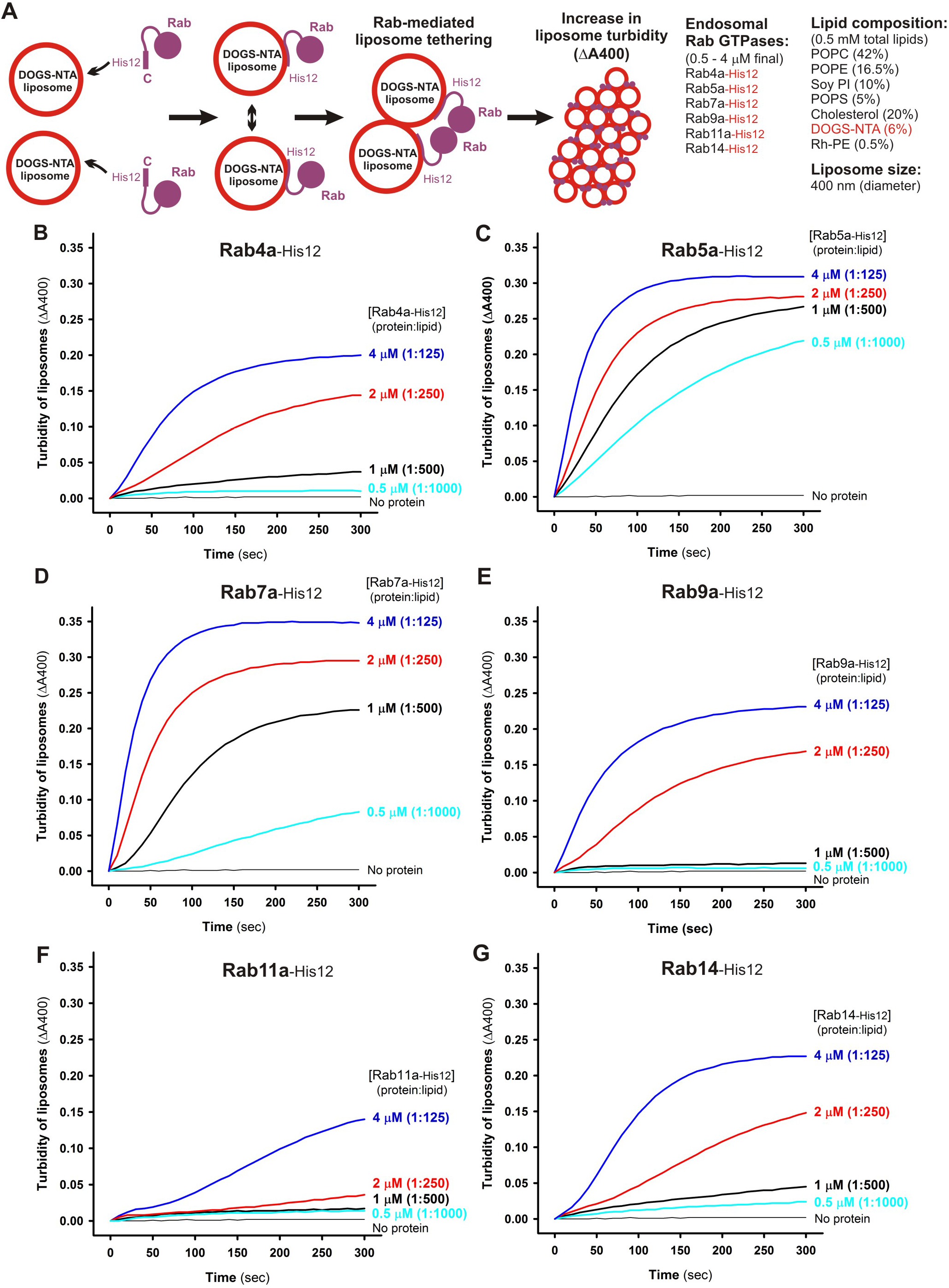
Endosomal Rab GTPases directly initiate membrane tethering by themselves in a chemically defined reconstitution system. (**A**) Schematic representation of liposome turbidity assays for testing Rab-mediated liposome tethering in **B-G**. (**B-G**) Endosomal Rab-His12 proteins (0.5 – 4 μM each), Rab4a-His12 (**B**), Rab5a-His12 (**C**), Rab7a-His12 (**D**), Rab9a-His12 (**E**), Rab11a-His12 (**F**), and Rab14-His12 (**G**), were incubated with synthetic liposomes bearing physiological mimic lipid composition (400 nm diameter; 0.5 mM lipids) in RB150 containing MgCl_2_ (5 mM) and DTT (1 mM) at room temperature for 300 sec. During incubation, turbidity changes of the Rab-liposome mixed reactions were monitored by measuring the absorbance at 400 nm. The protein-to-lipid molar ratios used for these turbidity reactions were from 1:1000 to 1:125, as indicated.

### Dissecting the membrane tethering potency of human Rab GTPases in a chemically defined reconstitution system

The intrinsic potency of human Rab GTPases to directly drive membrane tethering was thoroughly evaluated by the kinetics of increase in turbidity of liposome suspensions in the presence of Rab proteins, which can be monitored by measuring the absorbance at 400 nm (Fig. 2; Fig. 3) (18, 25–27). Rab-anchored liposomes were generated with purified Rab-His12 proteins and synthetic liposomes (400 nm in diameter) bearing a DOGS-NTA lipid and five major lipid species of phosphatidylcholine (PC), phosphatidylethanolamine (PE), phosphatidylinositol (PI), phosphatidylserine (PS), and cholesterol, roughly recapitulating lipid compositions of subcellular organelle membranes in mammalian cells (Fig. 2A) (28). Since the efficacy of Rab-mediated liposome tethering was dependent on lipid composition, liposome size, and total lipid concentration (supplemental Fig. S1) (18), we optimized the experimental conditions of liposome suspensions used for turbidity assays (Fig. 2A). With respect to Rab proteins tested in the tethering assays, as prior studies using the chemically defined reconstitution system had reported the intrinsic tethering activity of several Rabs in the endocytic trafficking pathways of yeast (the Rab5 ortholog Vps21p) (16) and human cells (Rab5a and Rab7a) (18), first we selected six representative endosomal Rabs (Rab4a, Rab5a, Rab7a, Rab9a, Rab11a, and Rab14) in human for comprehensively analyzing the tethering potency of Rab-family GTPases (Fig. 2B-G). By applying the liposome turbidity assay to the six Rab proteins at variable concentrations, ranging from 0.5 μM to 4 μM towards 0.5 mM total lipids in the reactions (Fig. 2A), rapid increase in liposome turbidity was triggered specifically either by Rab5a or Rab7a even at the low protein concentrations (0.5–1.0 μM, corresponding to the protein-to-lipid molar ratios of 1:1000–1:500) (Fig. 2C,D). This establishes the very high potency of these two endosomal Rabs to directly catalyze membrane tethering reactions *in vitro*, consistent with our prior study on membrane tethering mediated by human Rabs (18). Moreover, when assayed with the high protein concentrations of Rabs such as 4 μM (protein-to-lipid ratio, 1:125), we revealed that not only Rab5a and Rab7a (Fig. 2C,D) but also all of the other endosomal Rabs (Rab4a, Rab9a, Rab11a, and Rab14) retained the significant intrinsic capacity to initiate efficient tethering of synthetic liposomes by themselves (Fig. 2B,E,F,G). These data led us to propose the conserved function for all the endosomal Rab-family GTPases to physically link two distinct lipid bilayers together for membrane tethering events. It is also noteworthy that the observed Rab-mediated membrane tethering reactions were driven by homotypic Rab-Rab pairing specifically, but not by promiscuous Rab-Rab interactions (supplemental Fig. S2).

**Figure 3.**
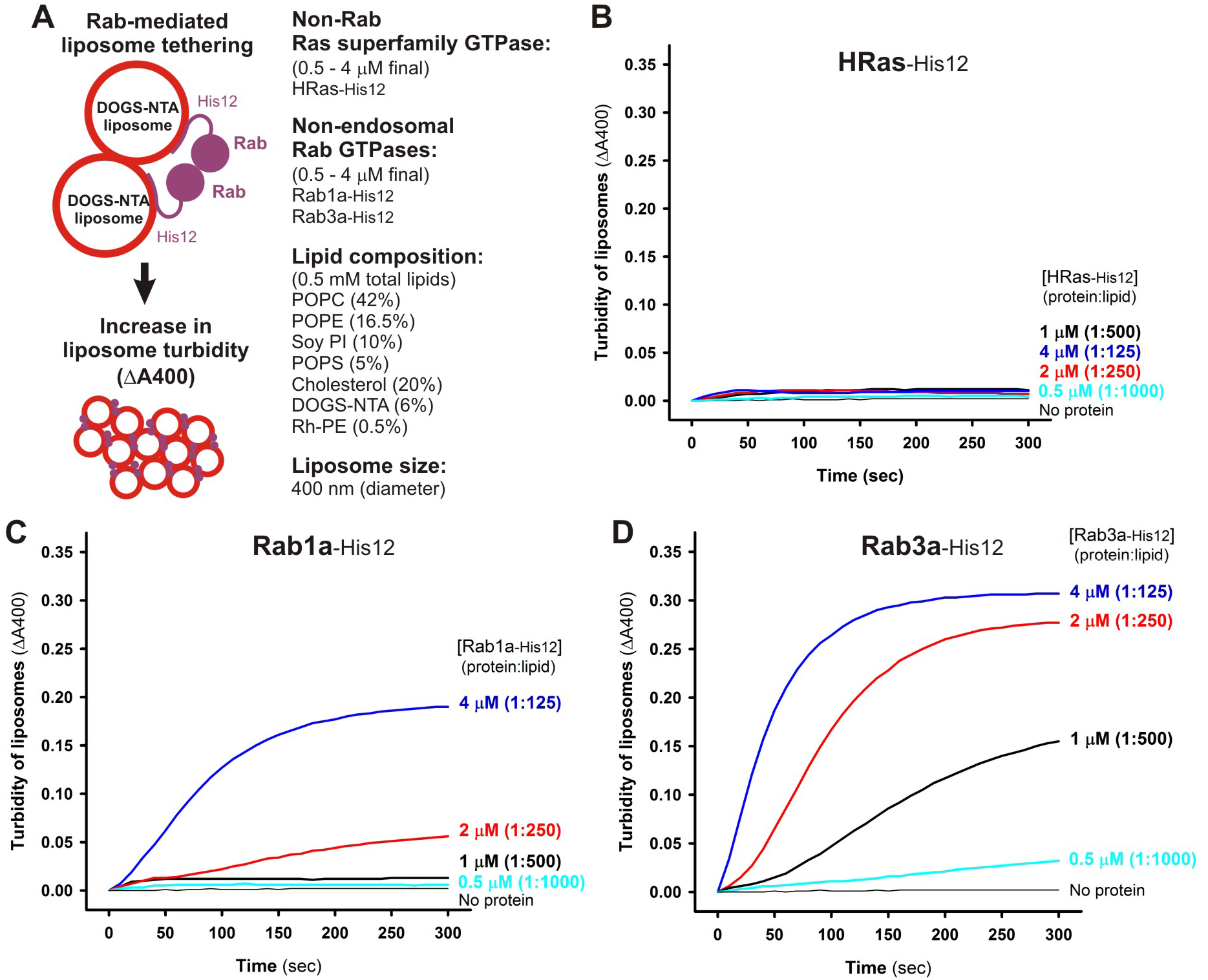
Non-endosomal Rab GTPases have the inherent potency to specifically mediate membrane tethering. (**A**) Schematic representation of liposome turbidity assays for the non-Rab Ras superfamily GTPase, HRas, and non-endosomal Rab GTPases, Rab1a in ER-Golgi traffic and Rab3a in exocytosis, in **B-D**. (**B-D**) Liposome turbidity assays were employed as in Fig. 2B-G, with HRas-His12 (**B**), Rab1a-His12 (**C**), and Rab3a-His12 (**D**) proteins (0.5 – 4 μM each in final) and physiological mimic synthetic liposomes (0.5 mM total lipids in final). The protein-to-lipid molar ratios used were indicated.

To what extent can the Rab protein densities on a membrane surface used in the current liposome turbidity assays (Fig. 2) recapitulate the physiological conditions at subcellular membranes in mammalian cells? Using synaptic vesicles as a model of a trafficking organelle, previous comprehensive proteomic and lipidomic analyses of highly purified synaptic vesicles from rat brain determined the average copy numbers per vesicle of the major protein constituents including Rab-family GTPases (29). Those quantitative data indicate that, on average, approximately 10 copies of Rab3a and 15 copies of the other Rab GTPases, thus 25 copies of Rab proteins in total are present in each single purified synaptic vesicle (29). Assuming (i) this copy number of synaptic Rabs (25 copies/vesicle), (ii) a mean outer diameter of synaptic vesicles of 42 nm (29), (iii) a typical bilayer thickness of 4 nm (30), and (iv) an average area occupied by a single phospholipid molecule of 0.65 nm^2^ (30), we estimated the surface areas of the outer and inner lipid layers of synaptic vesicles (5,540 nm^2^ and 3,630 nm^2^, respectively) that bore 14,100 lipid molecules in total, thereby giving approximately the Rab protein-to-lipid molar ratio of 1:560 (25 Rabs/vesicle, 14,100 lipids/vesicle). It should be noted that, in the current chemically defined system, Rab5a and Rab7a (at least) can trigger rapid and efficient membrane tethering at the Rab-to-lipid molar ratio of 1:500 (Fig. 2C,D), which is almost identical to the physiological ratio calculated above (Rab-to-lipid, 1:560). In addition to the Rab-to-lipid molar ratios, we also quantitatively estimated the membrane surface areas occupied by Rab proteins in the current reconstitution experiments (Fig. 2). Assuming that (i) Rab proteins are typically a spherical 25 kDa protein with an average radius of 2.0 nm (31), (ii) all of the Rab-His12 molecules added to the reactions are stably attached to liposomal membranes containing a DOGS-NTA lipid (Fig. 2A), and (iii) a single 400-nm-diameter liposome contains 1,520,000 lipid molecules (the total surface area, 985,000 nm^2^; the surface area/lipid, 0.65 nm^2^) (30), the percentages of the membrane surface coverage by Rab proteins were calculated to be in the range of 3.79% to 30.3% (0.5 μM – 4.0 μM Rab proteins; the Rab-to-lipid ratio, 1:1000 – 1:125) (Fig. 2). These estimated values of the surface coverage by Rab proteins reflect that Rab proteins occupy only a minor portion of the membrane surface areas of liposomes in the turbidity assays and thus appear to have plenty of space to interact and cooperate with other proteins and lipids on the membranes. Taken together, the Rab-to-lipid molar ratios and membrane surface areas occupied by Rabs that we estimated above support the idea that endosomal Rab proteins can drive rapid and efficient membrane tethering *in vitro* under the physiological or physiologically relevant conditions mimicking subcellular membranes in mammalian cells.

Our data of the current liposome turbidity assays have revealed the conserved membrane tethering potency of endosomal Rab proteins in the context of a physiologically relevant function (Fig. 2B-G). Next, we asked whether the intrinsic membrane tethering potency of human Rabs is the specialized function exclusively for Rab-family small GTPase proteins among the Ras superfamily GTPases (24) and, moreover, for the Rab proteins that are specifically localized at the endosomal compartments. To address this, we performed the turbidity assays for the HRas protein as the model of a non-Rab Ras-superfamily GTPase (Fig. 3A,B) and for two of the well-studied non-endosomal Rab proteins, Rab1a in ER-to-Golgi traffic and Rab3a in exocytosis (Fig. 3A,C,D), with the same range of the protein concentrations as tested in Fig. 2B-G (0.5 μM – 4.0 μM HRas or Rab proteins; the protein-to-lipid molar ratios, 1:1000 – 1:125).

Indeed, HRas had little membrane tethering potency, giving no significant increase in the turbidity of liposomes when assayed even at the highest HRas protein concentrations of 4.0 μM (HRas-to-lipid, 1:125) (Fig. 3B). Nevertheless, both of the non-endosomal Rab GTPases tested, Rab1a and Rab3a, retained the intrinsic capacity to directly induce efficient membrane tethering of synthetic liposomes (Fig. 3C,D), as shown earlier with six endosomal Rabs in Fig. 2. It should be noted that the difference of the *in vitro* membrane tethering potency between different Rab-family members and also between Rab proteins and HRas is not simply relied on their attachment to liposomal membranes in the tethering reactions (Fig. 4). To examine the membrane association of Rab and HRas proteins in the current systems, liposome co-sedimentation assays were employed (Fig. 4A) with the same experimental conditions as in the liposome turbidity assays (Fig. 2; Fig. 3; protein-to-lipid, 1:500), demonstrating that all the tested Rab-family and HRas proteins were able to comparably and stably bind to liposomal membranes (Fig. 4B-J). Thus, our data from the turbidity assays (Fig. 2; Fig. 3) suggest that the intrinsic potency to physically tether two apposed membranes is selectively encoded in the Rab-family members among the Ras-superfamily GTPases and likely to be fully conserved through all the Rab-family members functioning in the endocytic and secretory pathways.

**Figure 4.**
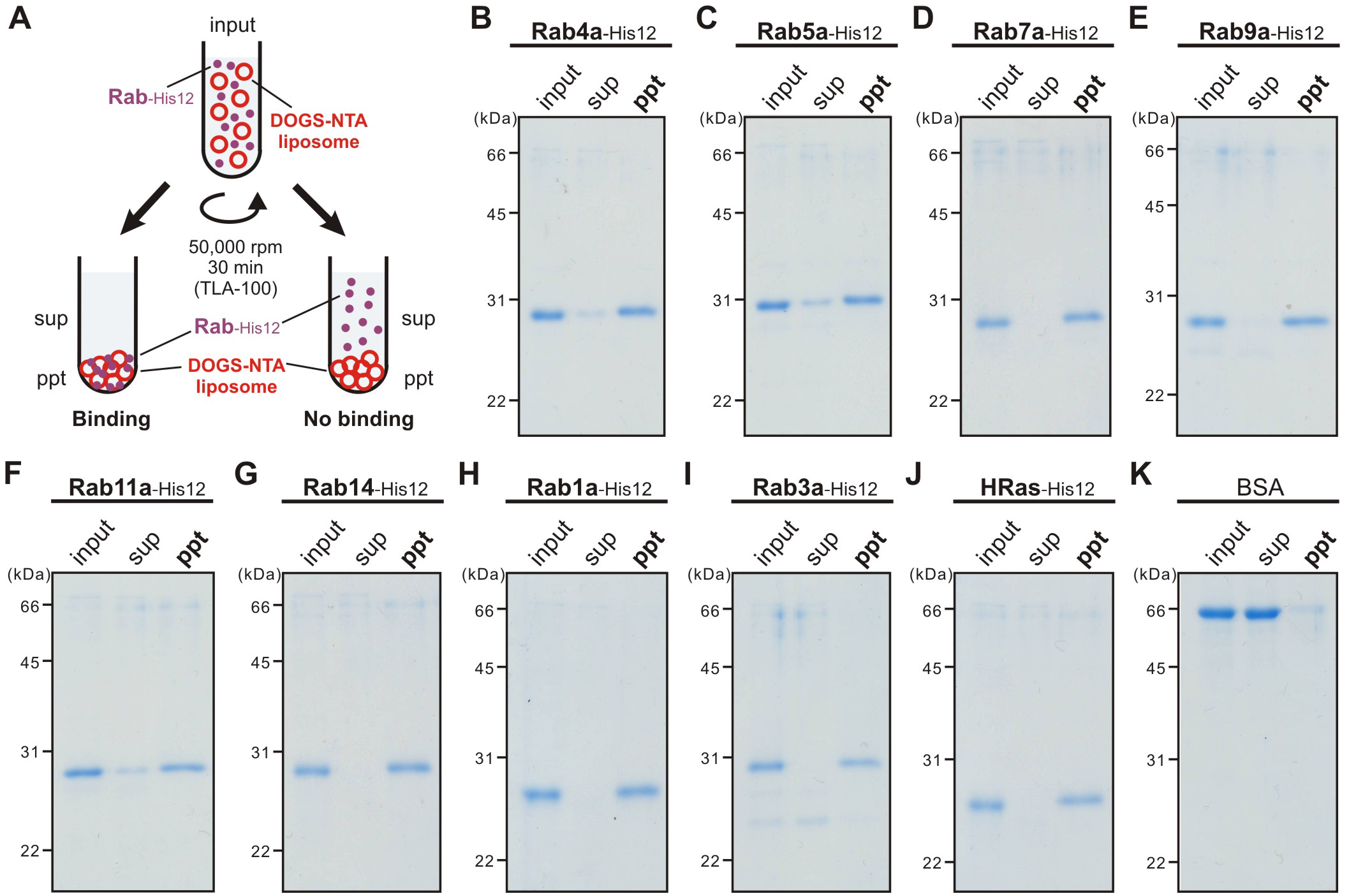
Membrane association of Rab-His12 proteins onto DOGS-NTA-bearing liposomes. (**A**) Schematic representation of liposome co-sedimentation assays for testing membrane attachment of Rab-His12 proteins used in Fig. 2 and Fig. 3. (**B-K**) Rh-labeled liposomes (400 nm diameter; 1 mM total lipids in final) were incubated (30°C, 30 min) with Rab4a-His12 (**B**), Rab5a-His12 (**C**), Rab7a-His12 (**D**), Rab9a-His12(**E**), Rab11a-His12 (**F**), Rab14-His12 (**G**), Rab1a-His12 (**H**), Rab3a-His12 (**I**), HRas-His12 (**J**), and BSA for a negative control (**K**) (2 μM final for each), and ultracentrifuged (50,000 rpm, 30 min, 4°C). The supernatants (sup) and precipitates (ppt) obtained were analyzed by SDS-PAGE and Coomassie Blue staining.

To further confirm the intrinsic membrane tethering capacity of Rab-family GTPases as their genuine function, we employed fluorescence microscopic observations of the reconstituted reactions of Rab-mediated membrane tethering (Fig. 5). The tethering reactions were incubated with rhodamine (Rh)-labeled fluorescent liposomes (0.5 mM lipids; 1000 nm diameter) and Rab-family or HRas proteins (4 μM proteins; protein-to-lipid, 1:125) (Fig. 5A) as in the liposome turbidity assays (Fig. 2; Fig. 3). Strikingly, whereas only small particles of non-tethered liposomes were observed when incubated in the absence of any Rab proteins (Fig. 5B,C) or with the control HRas protein (Fig. 5T,U), all of the tested endosomal and non-endosomal Rab proteins (Rab4a, Rab5a, Rab7a, Rab9a, Rab11a, Rab14, Rab1a, and Rab3a) were able to specifically induce the formation of substantial massive clusters of liposomes (Fig. 5D-S). Liposome clusters observed in the fluorescence images were then quantitatively analyzed for the particle size distributions (Fig. 6A), yielding the average sizes of Rab-mediated liposome clusters ranging from 7.8 μm^2^ to 55 μm^2^, which are much larger than those of the control reactions (0.92 μm^2^ and 0.80 μm^2^ for the reactions without any Rabs and with HRas, respectively) (Fig. 6B). These our morphological analyses of Rab-mediated membrane tethering reactions by fluorescence microscopy were fully consistent with the results obtained in the liposome turbidity assays (Fig. 2; Fig. 3), thus further establishing the working model that Rab-family proteins themselves can be a *bona fide* membrane tether to directly and physically link two distinct lipid bilayers of subcellular membranes in intracellular membrane tethering.

**Figure 5.**
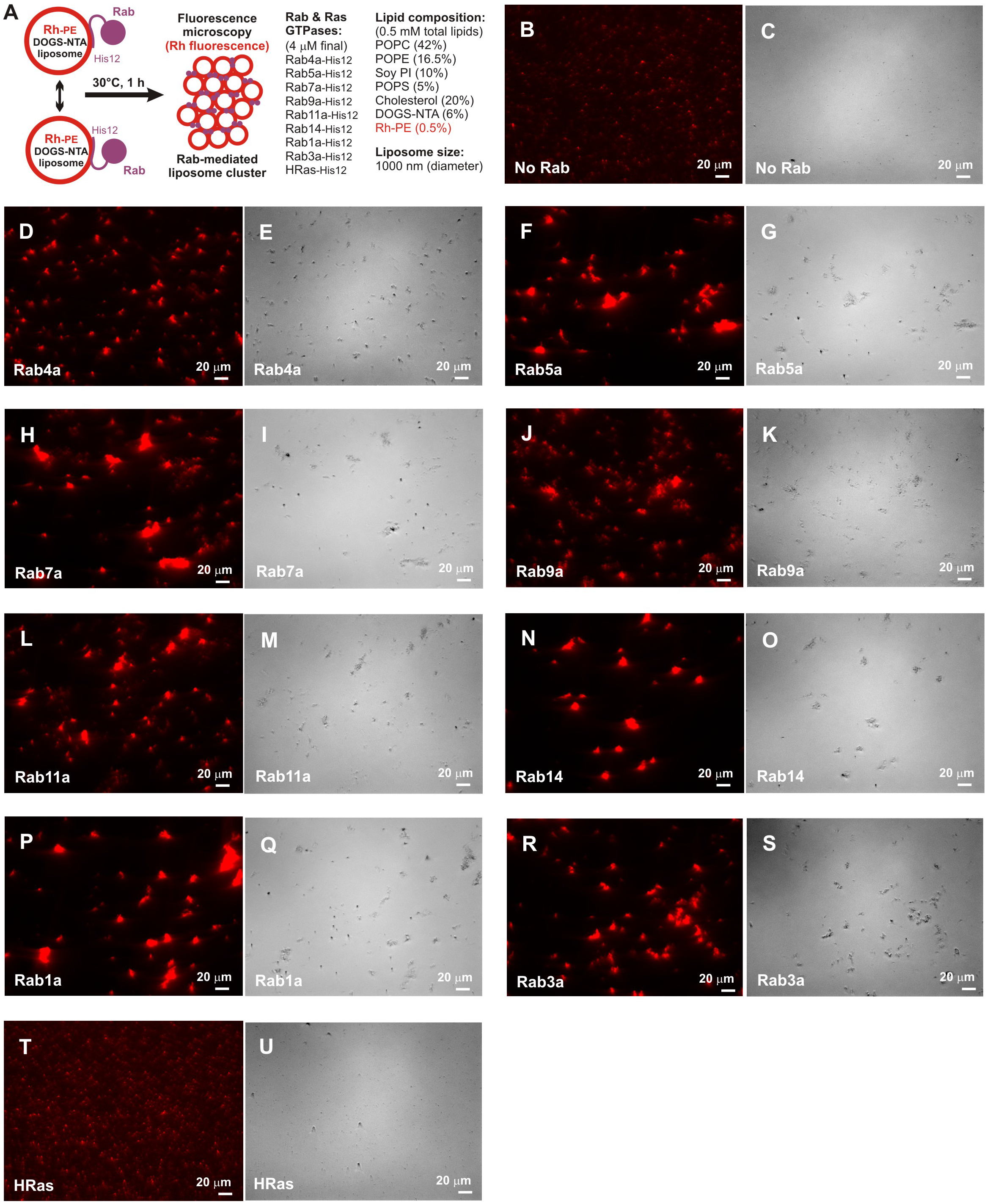
Rab-mediated membrane tethering induces the formation of massive liposome clusters. (**A**) Schematic representation of fluorescence microscopic observations of Rab-mediated liposome clusters. (**B-U**) Fluorescence images (**B, D, F, H, J, L, N, P, R, T**) and bright field images (**C, E, G, I, K, M, O, Q, S, U**) of Rab-mediated liposome clusters. Fluorescently-labeled liposomes bearing Rh-PE (1000 nm diameter; 0.5 mM lipids in final) were incubated at 30°C for 1 h, in the absence (**B, C**) and presence of the Rab- and Ras-family GTPases (4 μM each in final), including Rab4a-His12 (**D, E**), Rab5a-His12 (F, G), Rab7a-His12 (**H, I**), Rab9a-His12 (**J, K**), Rab11a-His12 (**L, M**), Rab14-His12 (**N, O**), Rab1a-His12 (**P, Q**), Rab3a-His12 (**R, S**), and HRas-His12 (**T, U**), and subjected to fluorescence microscopy. Scale bars: 20 μm.

**Figure 6.**
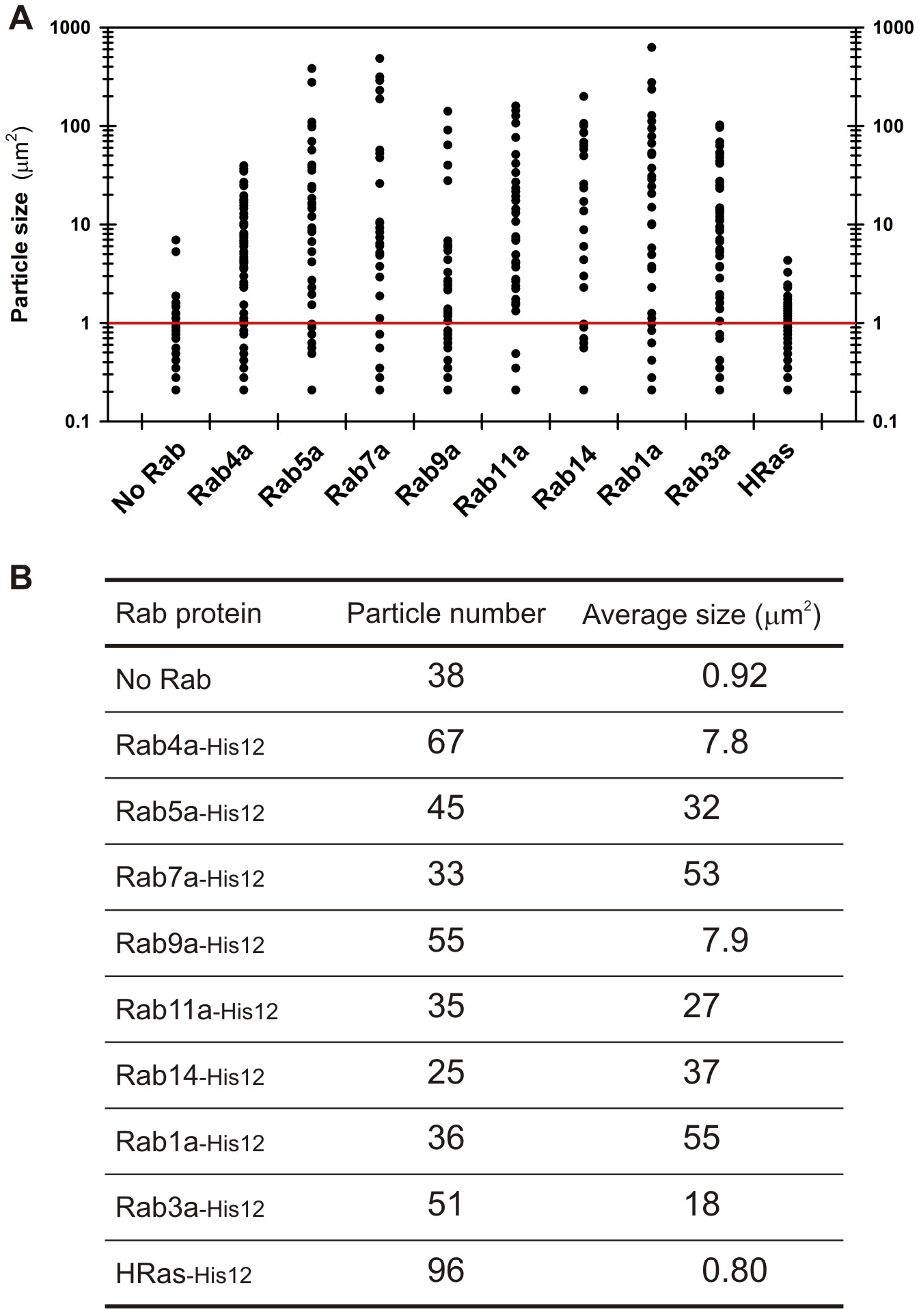
Particle size distributions of liposome clusters induced by Rab-mediated membrane tethering. (**A**) Particle sizes of the Rab-mediated liposome clusters observed in the fluorescence images of Fig. 5. (**B**) Particle numbers and average sizes of the Rab-mediated liposome clusters observed in the fluorescence images of Fig. 5.

### Class V myosins, the Rab11a effectors, strongly and selectively promote membrane tethering mediated by the cognate Rab GTPase

By comprehensively and quantitatively characterizing *in vitro* membrane tethering reactions reconstituted with chemically defined lipid bilayers and purified recombinant proteins of eight Rab GTPases in human (Rab1a, Rab3a, Rab4a, Rab5a, Rab7a, Rab9a, Rab11a, and Rab14) (Figs. 1–6), we now establish that Rab-family proteins genuinely have the intrinsic potency to directly and specifically tether two distinct lipid bilayers together in the context of a physiologically relevant function. Nevertheless, it is noteworthy that there appears to be wide variability in the tethering capacity of human Rab-family GTPases, although Rab proteins share the highly conserved amino acid sequences and tertiary structures for their Ras-superfamily GTPase domains (Fig. 1A) (24, 32), which exhibited comparable GTP-hydrolysis activities for all the eight Rabs tested (Fig. 1C). For typical instances of variability in the Rab-mediated tethering potency, Rab5a and Rab7a were able to trigger rapid membrane tethering at the protein-to-lipid molar ratios of 1:500 (Fig. 2C,D), whereas most of the other Rabs (Rab1a, Rab4a, Rab9a, Rab11a, and Rab14) showed little or no tethering activity under the same conditions of the Rab density on the membrane surface (Rab-to-lipid, 1:500) (Fig. 2B,E-G; Fig. 3C). These results led us to hypothesize that (i) the intrinsic tethering activity of Rab proteins is negatively autoregulated in general, especially in the case of the Rabs that cause very slow and inefficient membrane tethering by themselves (Rab1a, Rab4a, Rab9a, Rab11a, and Rab14) and thus (ii) specific Rab-interacting proteins (Rab effectors) drastically enhance the capacity of their cognate Rabs to drive membrane tethering. To test this hypothesis, we next attempted to reconstitute membrane tethering reactions with Rab11a, which had exhibited the lowest tethering potency among the eight Rabs tested in the earlier liposome turbidity assays (Fig. 2F), and the cognate Rab11a effectors, class V myosin motor proteins (Myo5A and Myo5B) (Fig. 7). Although any Rab11a effectors identified, except for the Exoc6/Sec15 exocyst subunit (33–35), have never been proposed or reported to be directly involved in membrane tethering events (4, 5, 7), it has been thought that, as Rab effectors, class V myosins directly bind and cooperate with the cognate Rab11a on transport vesicles to regulate the specificity of membrane trafficking (36–39).

**Figure 7.**
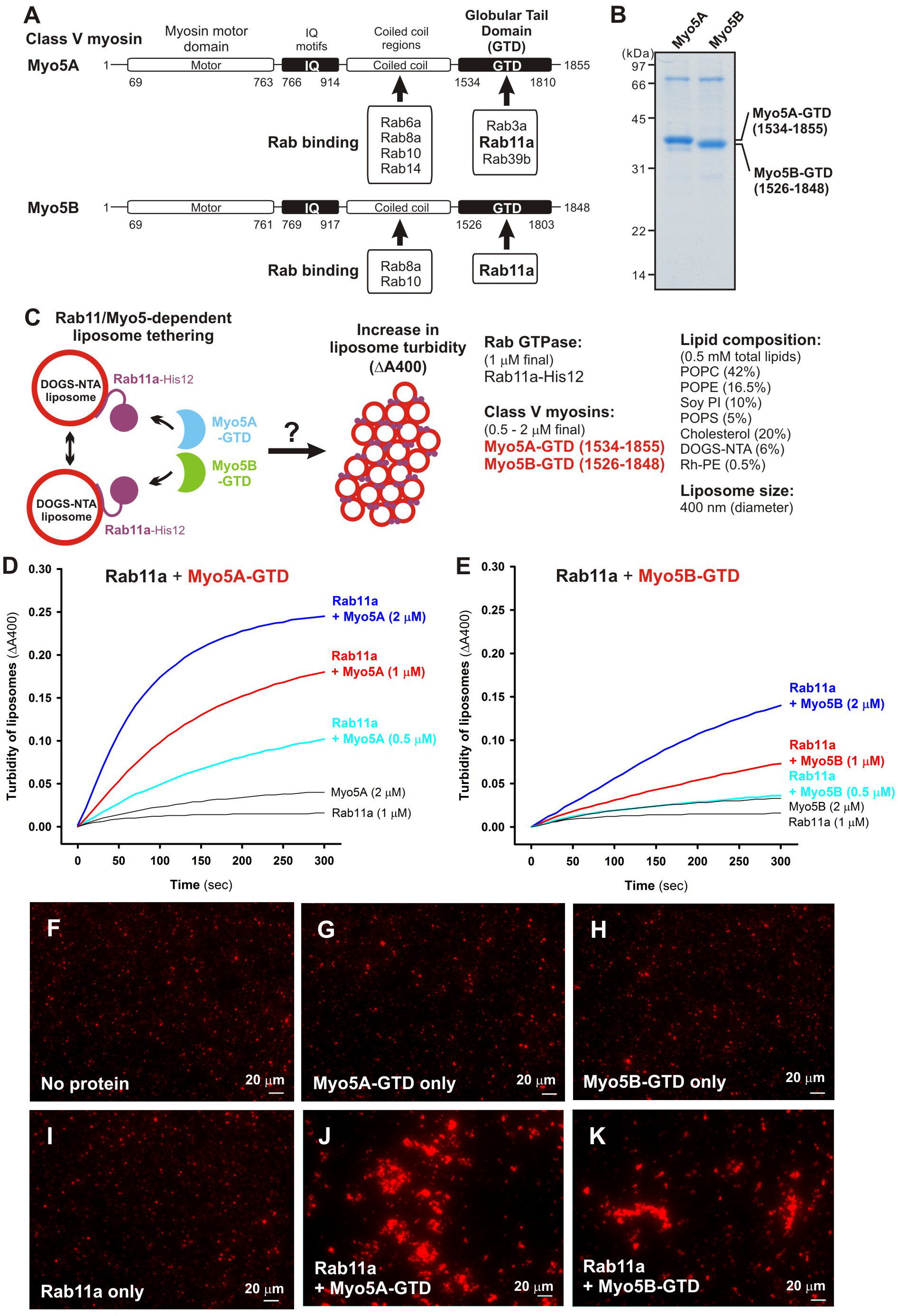
Class V myosin globular tail domains, Myo5A-GTD and Myo5B-GTD, strongly stimulate Rab11a-dependent membrane tethering. (**A**) Schematic representation of class V myosins in human, Myo5A and Myo5B, showing their amino acid residues and domains including myosin motor domains, IQ motifs, coiled coil regions, and globular tail domains (GTDs). Representative Myo5-interacting Rab GTPases and the Rab-binding regions in Myo5A and Myo5B are indicated. (**B**) The Coomassie Blue-stained gel of purified Myo5A-GTD and Myo5B-GTD proteins, which are comprised of the amino acid residues 1534-1855 and 1526-1848, respectively. (**C**) Schematic representation of liposome turbidity assays for testing Rab11a- and Myo5-GTD-dependent liposome tethering in **D, E**. (**D, E**) Liposome turbidity assays were employed with Rab11a-His12 (1 μM final) as in Fig. 2F, but in the presence of Myo5A-GTD (**D**) and Myo5B-GTD (**E**) (0.5 – 2 μM final). (**F-K**) Fluorescence images of Rab11a-mediated liposome clusters in the presence of Myo5-GTDs. Rab11a-His12 (3 μM final) and Myo5A-GTD or Myo5B-GTD (3 μM final) were preincubated at 30°C for 30 min, mixed with Rh-labeled liposomes (1000 nm diameter; 0.8 mM lipids in final), further incubated (30°C, 30 min), and subjected to fluorescence microscopy (**J, K**). For a control, Rab11a-His12, Myo5-GTD, or both were omitted from the reactions where indicated (**F-I**). Scale bars: 20 μm.

To reconstitute the Rab11a- and class V myosin-dependent membrane tethering reaction in a chemically defined system, we purified the globular tail domains of class V myosin proteins (Myo5A-GTD and Myo5B-GTD) in human, which correspond to the C-terminal residues 1534-1855 of myosin 5a (Myo5A) and the residues 1526-1848 of myosin 5b (Myo5B) (Fig. 7A,B). It should be noted that previous biochemical and structural studies indicated that these globular tail domains of Myo5A and Myo5B were necessary and sufficient for binding to Rab11a (Fig. 7A) (39–42). Using purified Myo5A-GTD and Myo5B-GTD proteins (Fig. 7B), we employed liposome turbidity assays to test whether class V myosin proteins have an effect on membrane tethering reactions mediated by the cognate Rab11a (Fig. 7C-E). Strikingly, Rab11a-mediated tethering was strongly promoted by addition of an equimolar amount (1 μM) or a 2-fold molar excess (2 μM) of Myo5A-GTD (Fig. 7D) or Myo5B-GTD (Fig. 7E) over Rab11a (1 μM; Rab-to-lipid ratio, 1:500), exhibiting much higher initial rates of the turbidity increase than those in the absence of Myo5-GTD proteins. We also showed that a non-interacting control protein (BSA) had no stimulatory effect on the tethering activity of Rab11a (supplemental Fig. S3). It should be noted that the molar ratio of the yeast ortholog of class V myosins (Myo2p) to its cognate Rab protein (Sec4p) in yeast cells was estimated to be about an equimolar ratio using quantitative immunoblots (43), indicating that the Myo5-to-Rab molar ratios used in the current reconstitution assays (Fig. 7C-E) are not far from those of physiological conditions. Morphological changes of liposomes in the tethering reactions with Rab11a and Myo5-GTDs were also analyzed by fluorescence microscopy (Fig. 7F-K), indicating that Rab11a and Myo5-GTD proteins synergistically and specifically induced the formation of massive clusters of liposomes (Fig. 7J,K). Thus, the current results from these two independent assays in a chemically defined system uncover the novel function of class V myosins to directly support membrane tethering mediated by their cognate Rab11a GTPase.

Since class V myosins are recognized as Rab effectors that, in general, selectively interact with the GTP-bound from of Rab proteins (38), we next tested the guanine nucleotide dependence of Rab11a- and Myo5-GTD-mediated membrane tethering by employing the liposome turbidity assays (Fig. 8A). Indeed, the addition of GTP (1 mM) drastically stimulated the synergistic action of Rab11a and Myo5A-GTD (Fig. 8B) or Myo5B-GTD (Fig. 8C) upon driving rapid membrane tethering, but the presence of GTP had no effect on the tethering reactions bearing either Rab11a or Myo5-GTD proteins alone (Fig. 8B,C). The GTP independence of membrane tethering mediated by Rab11a alone is consistent with our previous data with Rab5a and the other Rab isoforms (18). Considering that the targeting of Rab proteins to subcellular membrane compartments is mediated by GDP/GTP exchange and Rab guanine nucleotide exchange factors (44–46), these results suggest that the stable attachment of Rab-His12 proteins on DOGS-NTA-bearing membranes in the current system can bypass the GTP requirement for Rab-only tethering reactions (Fig. 8B,C) (18). The direct stimulation by GTP on Rab11a/Myo5-GTD-dependent membrane tethering was shown as a very specific effect when added the variable GTP concentrations ranging from 1 μM to 1 mM (Fig. 8D,E) and assayed with the other guanine nucleotides, GDP and GTPγS (Fig. 8F,G). Our data therefore reflect that Myo5-GTD proteins greatly enhance the tethering activity of Rab11a in a GTP-dependent manner, suggesting that the proteinprotein interactions of Myo5-GTD proteins with membrane-bound Rab11a-GTP proteins are required for facilitating membrane tethering. To experimentally test the association of Myo5-GTD proteins with Rab11a proteins on liposomal membranes, liposome co-sedimentation assays were performed under the typical conditions used for the turbidity assays in Fig. 8 (Rab-to-lipid ratio, 1:500; Rab-to-Myo5-GTD ratio, 1:1; 1 mM GTP) (Fig. 9A). However, we found that the amounts of Myo5A-GTD (Fig. 9B) and Myo5B-GTD (Fig. 9E) bound to and co-isolated with Rab11a-attached liposomes were comparable to those of the control reactions with protein-free liposomes in the absence of Rab11a-His12 (Fig. 9C,F) or in the presence of untagged Rab11a lacking a His12 tag (Fig. 9D,G). This indicates, unexpectedly, that the membrane association of Myo5-GTD proteins was not significantly promoted by the recruitment of Rab11a on a liposomal membrane surface. Considering that rapid and efficient Rab11a-mediated membrane tethering is definitely relied on the presence of Myo5-GTD proteins and GTP in the current reconstitution system (Fig. 7; Fig. 8), our results from the co-sedimentation assays (Fig. 9) may imply that a specific, but not a stable, interaction is required for the tethering reaction. Myo5-GTD in this assay appears to act as a “catalyst” of conformational change in Rab11a that is required for increased Rab-Rab interactions, and this conformational change may be additionally facilitated by GTP binding.

**Figure 8.**
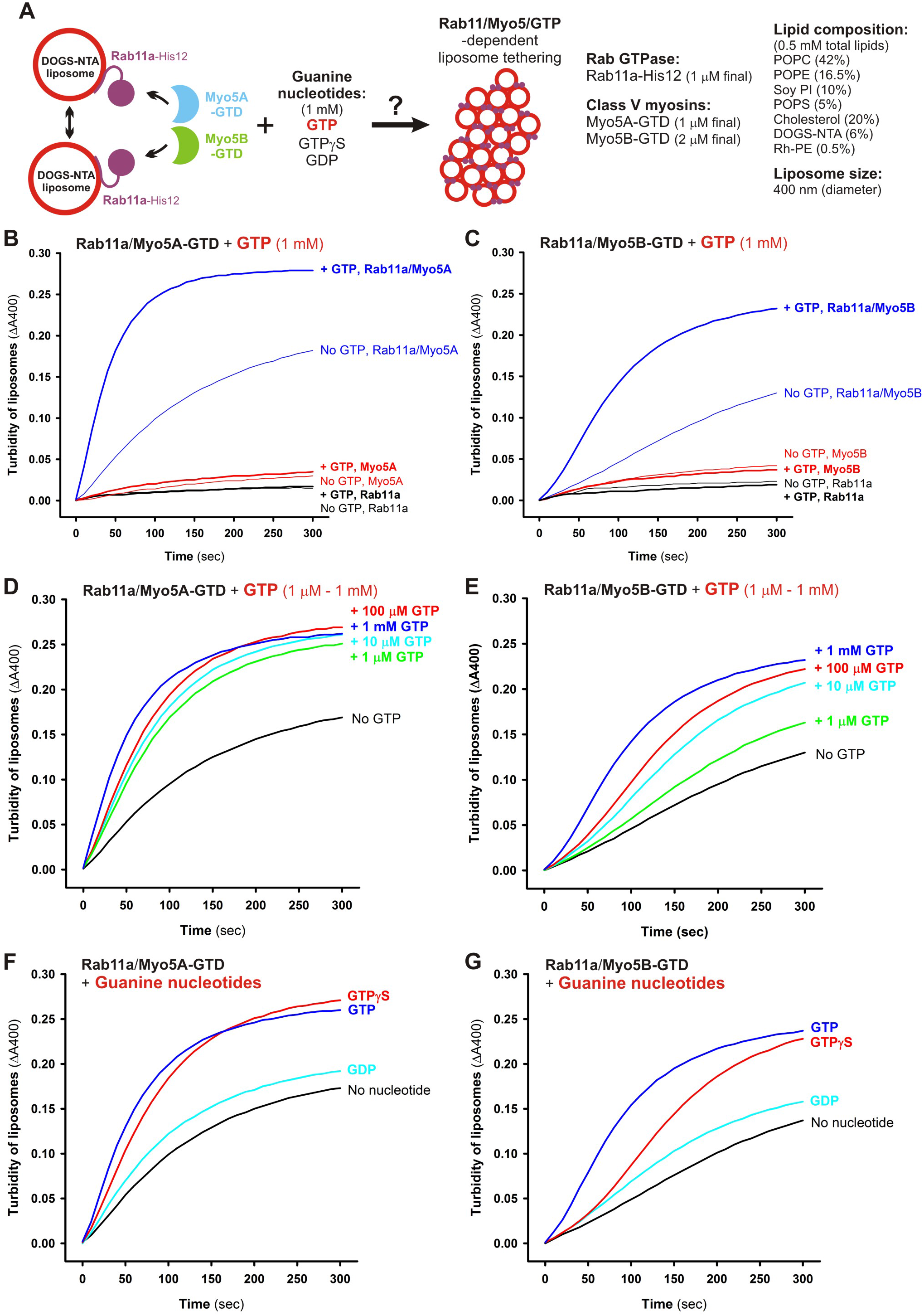
Guanine nucleotide dependence of Rab11a-mediated membrane tethering in the presence of Myo5A-GTD and Myo5B-GTD. (**A**) Schematic representation of liposome turbidity assays for testing Rab11a- and Myo5-GTD-dependent liposome tethering in the presence of GTP in **B-G**. (**B, C**) Rab11a/Myo5-dependent membrane tethering is strongly and specifically promoted by the addition of GTP. Liposome turbidity assays with Rab11a-His12 (1 μM) and Myo5A-GTD (1 μM) (B) or Myo5B-GTD (2 μM) (**C**) were performed as in Fig. 7D, E, but in the presence of GTP (1 mM). (**D, E**) Liposome turbidity assays were employed with Rab11a-His12 and Myo5A-GTD (**D**) or Myo5B-GTD (**E**) as in **B, C**, in the presence of various concentrations of GTP (1 μM – 1 mM). (**F, G**) Liposome turbidity assays were employed with Rab11a-His12 and Myo5A-GTD (**F**) or Myo5B-GTD (**G**) as in **B, C**, in the presence of GTP, GTPγS, and GDP (1 mM for each).

**Figure 9.**
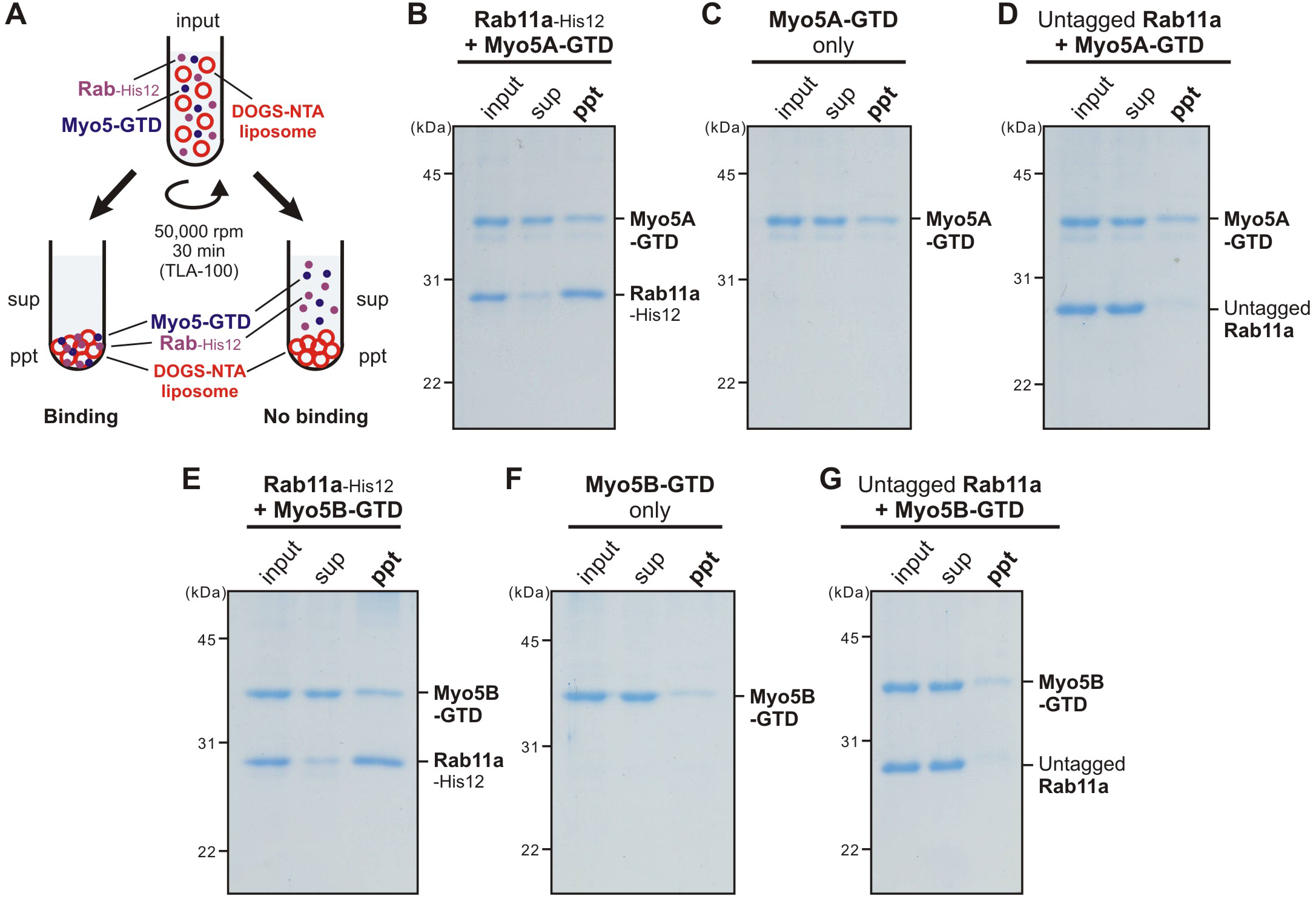
Membrane association of Myo5-GTD proteins onto Rab11a-anchored liposomes. (**A**) Schematic representation of liposome co-sedimentation assays for testing membrane binding of Myo5A-GTD and Myo5B-GTD to Rab11a-bound liposomes. (**B-G**) Liposome co-sedimentation assays were employed as in Fig. 4, with Rh-labeled liposomes (400 nm diameter; 1 mM lipids) and Rab11a-His12 (2 μM) (**B, E**), but in the presence of Myo5A-GTD (2 μM) (**B-D**), Myo5B-GTD (2 μM) (**E-G**), and GTP (1 mM). For a control, the reactions without Rab11a-His12 (**C, F**) or with the untagged form of Rab11a lacking a His12 tag (untagged Rab11a) (**D, G**) were also tested. The supernatants (sup) and precipitates (ppt) obtained were analyzed by SDS-PAGE and Coomassie Blue staining.

Our current studies have uncovered the novel function of class V myosins to directly promote membrane tethering mediated by their cognate Rab11a GTPase in a GTP-dependent manner, by thoroughly characterizing reconstituted membrane tethering in the chemically defined systems with purified Rab11a and Myo5-GTD proteins (Figs. 7–9). However, it could still be argued that Myo5-GTD proteins can non-physiologically induce tethering or aggregation of liposomal membranes coated by other noncognate Rab-His12 proteins or even coated by any types of soluble proteins modified with a polyhistidine tag. In this context, to more strictly validate the specificity of Rab11a/Myo5-GTD-dependent membrane tethering, we further explored the liposome turbidity assays in the presence of Myo5-GTD proteins (Fig. 10A), not only for the cognate Rab11a but also for the other four Rab proteins including Rab1a, Rab4a, Rab9a, and Rab14, which had shown relatively slow and inefficient membrane tethering by themselves, similar to Rab11a (Fig. 2; Fig. 3). Although Rab11a and either Myo5A-GTD or Myo5B-GTD proteins synergistically triggered very rapid and efficient membrane tethering (Fig. 10B,C), fully consistent with the earlier results in Fig. 8, these Myo5-GTD proteins indeed had only minor or little effect on the tethering potency of Rab1a, Rab4a, Rab9a, and Rab14 (Fig. 10B,C).

**Figure 10.**
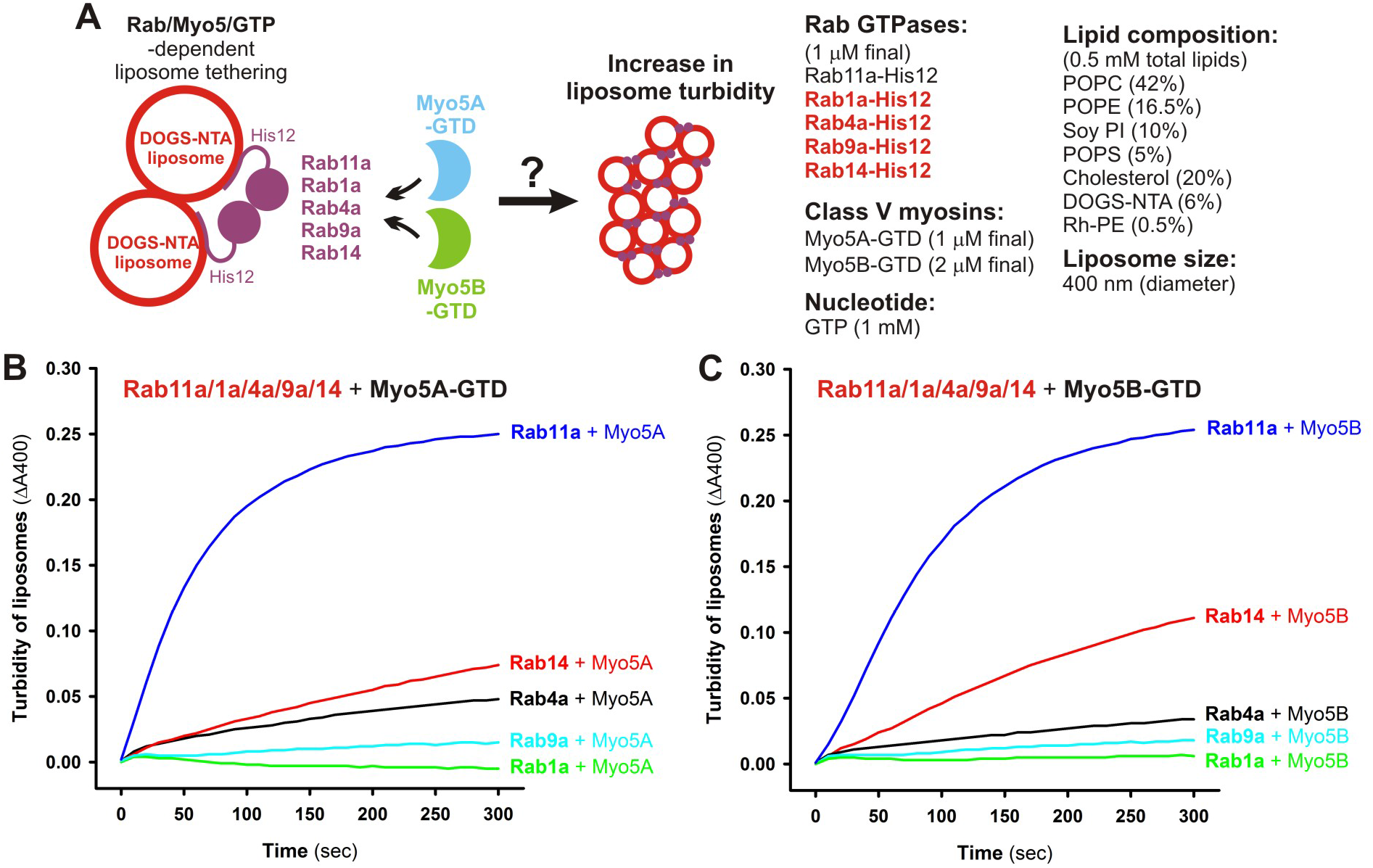
Myo5A-GTD and Myo5B-GTD selectively activate Rab11a-dependent membrane tethering. (**A**) Schematic representation of liposome turbidity assays in **B, C**, for the various Rab GTPases (Rab11a, Rab1a, Rab4a, Rab9a, and Rab14) in the presence of Myo5-GTDs and GTP. (**B, C**) Myo5-GTDs specifically promote efficient membrane tethering mediated by the cognate Rab GTPase, Rab11a. Liposome turbidity assays were employed with Myo5A-GTD (**B**) or Myo5B-GTD (**C**) and GTP, as in Fig. 8B, C, but for Rab11a, Rab1a, Rab4a, Rab9a, and Rab14 GTPases.

In addition to the Rab selectivity (Fig. 10), to further establish the specificity of the Rab11a-and Myo5-dependent membrane tethering, we next tested whether the membrane attachment of Rab11a on both of two opposing membranes is critical to catalyzing the tethering reactions, by employing the streptavidin bead-based liposome tethering assays (18, 52) (Fig. 11A-C) and fluorescence microscopy (Fig. 11D-G). In the streptavidin bead-based assays, purified proteins of Rab11a-His12 and Myo5-GTDs were incubated with two types of the DOGS-NTA liposomes bearing either biotin-PE/Rh-PE or FL-PE (fluorescein-PE) and a streptavidin-coated bead (Fig. 11A). Following the incubation of the reaction mixtures, FL fluorescence of the FL-PE liposomes co-isolated with the biotin-PE liposomes bound to the beads was measured, for quantitatively evaluating the membrane tethering capacities of these protein components (Fig. 11B,C). Consistent with the results of the liposome turbidity assays and fluorescence microscopic observations in Fig. 7, efficient membrane tethering was specifically triggered when added both Rab11a and Myo5-GTD proteins to the reactions (Fig. 11B, lanes 5 and 6). However, strikingly, the tethering potency of Rab11a and Myo5-GTDs was thoroughly abolished by omitting a DOGS-NTA lipid from one of the liposomes used, FL-PE liposomes (Fig. 11C, lanes 2 and 4). This reflects that Rab11a needs to be anchored on both of two opposing membrane surfaces for initiating membrane tethering in the presence of Myo5-GTDs. We then addressed the same issue, the requirement of membrane-bound Rab11a for tethering, by fluorescence microscopy with the Rh- and FL-labeled liposomes (Fig. 11D-G). As expected from the results of the bead-based tethering assays (Fig. 11B,C), addition of Rab11a and Myo5-GTDs to the liposome suspensions was able to induce the formation of massive liposome clusters bearing both of the Rh-PE liposomes and FL-PE liposomes (Fig. 11D,F). Nevertheless, when tested with the Rh-PE liposomes lacking a DOGS-NTA lipid instead, Rab11a and Myo5-GTDs no longer had their capacities to support the stable association of the Rh-PE liposomes with the FL-PE liposome clusters (Fig. 11E,G). Therefore, these results from the two independent reconstitution assays clearly demonstrate that Rab11a- and Myo5-mediated membrane tethering requires the membrane attachment of Rab11a proteins on both of two opposing membranes destined to tether, thus establishing the assemblies of membrane-anchored Rab11a proteins in *trans* to physically link two distinct lipid bilayers in this tethering process.

**Figure 11.**
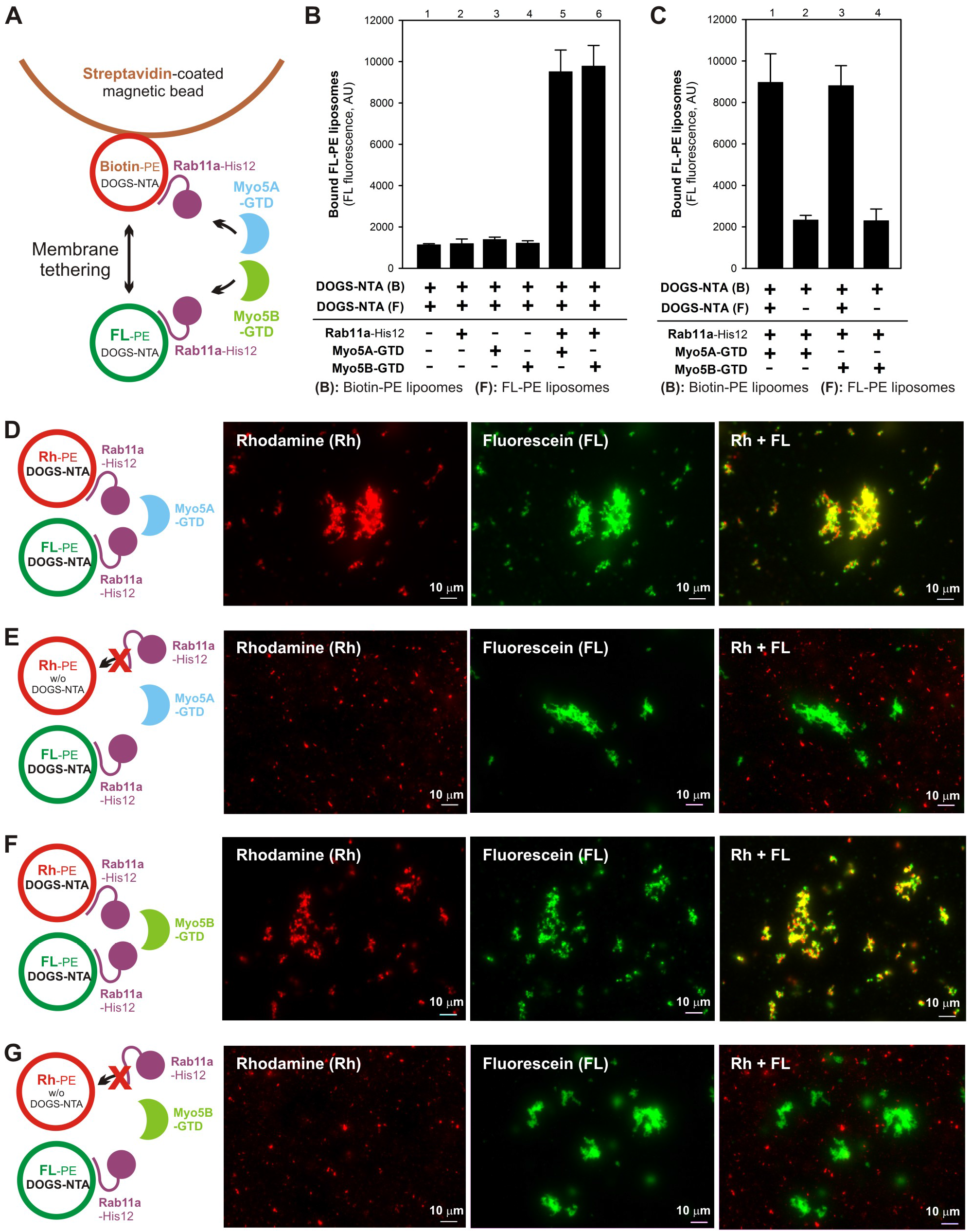
Rab11a- and Myo5-GTD-dependent membrane tethering requires the membrane attachment of Rab11a on both of two opposing membranes destined to tether. (**A**) Schematic representation of the streptavidin bead-based liposome tethering assay in **B, C**, using two types of liposomes bearing either biotin-PE/DOGS-NTA/Rh-PE or FL-PE/DOGS-NTA and purified Rab11a-His12 and Myo5-GTD proteins. (**B**) Streptavidin bead-based liposome tethering assays with Rab11a-His12 and Myo5-GTDs. The biotin-labeled liposomes and FL-labeled liposomes were incubated with streptavidin-coated magnetic beads (30°C, 2 hours), mixed with Rab11a-His12 and Myo5A-GTD or Myo5B-GTD, and further incubated (30°C, 10 min). The FL-labeled liposomes tethered to the biotin-labeled liposomes on streptavidin beads were quantified by measuring the FL fluorescence. For a control, Rab11a, Myo5A-GTD, and Myo5B-GTD were omitted from the reactions, where indicated (lanes 1-4). AU, arbitrary units. (**C**) Streptavidin bead-based liposome tethering assays were employed with Rab11a-His12 and Myo5-GTDs as in B, but the FL-labeled liposomes lacking DOGS-NTA were used instead where indicated (lanes 2 and 4). (**D-G**) Fluorescence microscopy was performed as in Fig. 7F-K, with Rh-labeled liposomes (0.4 mM lipids), FL-labeled liposomes (0.4 mM lipids), Rab11a-His12 (3 μM), and Myo5A-GTD (**D, E**) or Myo5B-GTD (**F, G**) (3 μM each). The Rh-labeled liposomes lacking DOGS-NTA were used instead where indicated (**E, G**). Scale bars: 10 μm.

Taken together, our data faithfully reflect that both Myo5A-GTD and Myo5B-GTD recognize and act upon exclusively the membrane-anchored form of the cognate Rab11a at the membrane surfaces of lipid bilayers, thereby selectively activating Rab11a-mediated membrane tethering reactions that are driven by *trans*-Rab protein assemblies. The current findings from reconstituted membrane tethering with purified Rab and Myo5-GTD proteins (Figs. 7–11) provide new insights into how Rab GTPases on subcellular membranes (transport vesicles, organelle membranes, or the plasma membrane) and class V myosin motors on actin cytoskeletal tracks cooperate with each other to promote membrane tethering and how their functions synergistically contribute to the spatiotemporal specificity of membrane tethering and beyond.

### Conclusions

Membrane tethering is one of the most critical and important processes to determine the specificity of membrane traffic in eukaryotic endomembrane systems, delivering the correct sets of cargoes (proteins, lipids, etc.) to the correct locations (organelles, the plasma membrane, the extracellular space, etc.) (8). From a large body of prior studies using genetic, cell biological, biochemical, and structural biological approaches on this essential membrane tethering process, we generally recognize that a number of diverse but specific types of “Rab effectors” have been described as the so-called “tethering factors”, such as coiled-coil tethering proteins and multisubunit tethering complexes (10, 47, 48). However, for the most cases, it remains ambiguous indeed whether these “tethering factors” identified are a *bona fide* membrane tether that physically links two distinct opposing membranes by itself or, furthermore, whether they are even involved directly in membrane tethering events (17). In this context, in order to thoroughly define and characterize the molecular functions in membrane tethering for Rab-family GTPases and class V myosin cytoskeletal motor proteins as the specific Rab effectors, we have undertaken to recapitulate the membrane tethering machinery in the chemically defined systems reconstituted with synthetic lipid bilayers of liposomes and purified proteins of human Rab GTPases and class V myosins (Fig. 1; Fig. 2; Fig. 7).

Our current *in vitro* reconstitution studies present three new findings supporting the working model of intracellular membrane tethering mediated by Rab-family GTPases as a genuine membrane tether: (i) All the eight Rab proteins tested, including both endosomal and non-endosomal Rabs, have the intrinsic capacity to physically tether two apposed membranes under the experimental conditions mimicking the lipid composition and Rab density of subcellular organelles/vesicles in mammalian cells, establishing the conserved tethering function of the Rab-family members (Fig. 2; Fig. 3; Fig. 5). (ii) The tethering capacity of Rab proteins is a specific function among the Ras superfamily GTPases and yet highly variable for each of different Rab proteins (Fig. 2; Fig. 3; Fig. 5). This suggests that the Rab subfamily-specific (RabSF) motifs could be the regions responsible for controlling the membrane tethering potency of each Rab (49, 50). Finally, (iii) the Rab effectors, class V myosins, can drastically and specifically promote membrane tethering mediated by their cognate Rab GTPase, Rab11a, in a GTP-dependent manner (Fig. 7; Fig. 8; Fig. 10; Fig. 11). This finding leads us to hypothesize that cytoskeletal motor proteins, including class V myosins and also microtubule-based motor proteins, act as “tethering factors” (not “tethers”) that directly and positively regulate Rab-mediated membrane tethering (37, 51). Understanding a detailed picture of the protein machinery of Rab-mediated membrane tethering will need further studies using a chemically defined reconstitution system, which may focus on testing tethering by the heterotypic Rab combinations, determining the specific regions in Rab molecules responsible for tethering, and reconstituting Rab-mediated tethering in the presence of different types of Rab effectors.

## Experimental procedures

### Protein expression and purification

The coding sequences for the eight isoforms of Rab-family GTPases (Rab1a, Rab3a, Rab4a, Rab5a, Rab7a, Rab9a, Rab11a, and Rab14) in human and for human HRas GTPase were amplified by polymerase chain reaction (PCR) with Human Universal QUICK-Clone cDNA II (Clontech, Mountain View, CA) as a template cDNA and KOD-Plus-Neo DNA polymerase (Toyobo, Osaka, Japan), as described (18). The amplified PCR fragments contained the sequence encoding a human rhinovirus (HRV) 3C protease site (Leu-Glu-Val-Leu-Phe-Gln-Gly-Pro) upstream of the initial ATG codons and the sequence encoding polyhistidine residues (His12) downstream of the codons for a C-terminal residue, yielding the full-length Rab and HRas proteins with three extra N-terminal residues (Gly-Pro-Gly) and a C-terminal His12 tag after HRV 3C protease cleavage. For Rab11a, the PCR fragment without the polyhistidine-coding sequence was also amplified as above, to prepare the Rab11a protein lacking the C-terminal His12 tag (untagged Rab11a). All of these PCR fragments for the Rab-family and HRas GTPases were inserted into a pET-41 Ek/LIC vector (Novagen, Madison, WI), which is designed to express an N-terminal GST-His6-tagged protein, by the ligation-independent cloning method (Novagen).

Recombinant Rab and HRas proteins were expressed in the *Escherichia coli* BL21(DE3) cells (Novagen) harboring the pET-41-based vectors in Lysogeny Broth (LB) medium (1 liter each) with kanamycin (50 μg/ml) by induction with 0.1 mM IPTG at 37°C for 3 hours. *E. coli* cells harvested after IPTG induction were resuspended in 40 ml each of RB150 (20 mM Hepes-NaOH, pH 7.4, 150 mM NaCl, 10% glycerol) containing 0.1 mM GTP, 5mM MgCl_2_, 1 mM DTT, 1 mM PMSF, and 1.0 μg/ml pepstatin A, freeze-thawed in a liquid nitrogen and a water bath at 30°C, lysed by sonication using UD-201 ultrasonic disrupter (Tomy Seiko, Tokyo, Japan), and then ultracentrifuged at 50,000 rpm for 75 min at 4°C with a 70 Ti rotor (Beckman Coulter, Indianapolis, IN). The supernatants obtained were mixed with COSMOGEL GST-Accept beads (50% slurry, 4 ml each; Nacalai Tesque, Kyoto, Japan) and were incubated at 4°C for 2 hours with gentle agitation to isolate GST-His6-tagged Rab-His12 and HRas-His12 proteins. The protein-bound GST-Accept beads were washed four times in RB150 containing 5 mM MgCl_2_ and 1 mM DTT (8 ml each), resuspended in the same buffer (4 ml each), supplemented with HRV 3C protease (4 units/ml final; Novagen), and incubated without agitation (4°C, 16 hours) to cleave off and elute Rab-His12 and HRas-His12 proteins. After centrifugation of the bead suspensions (15,300 g, 10 min, 4°C), purified Rab-His12 and HRas-His12 proteins were harvested from the supernatants.

For class V myosins, the coding sequences of the globular tail domains of human myosin 5a (Myo5A-GTD; residues 1534-1855) and myosin 5b (Myo5B-GTD; residues 1526-1848) with the sequence encoding a HRV 3C-protease site upstream of the initial codons were amplified by PCR as above and then cloned into a pET-30 Ek/LIC vector (Novagen) expressing an N-terminal His6-tagged protein. Recombinant Myo5A-GTD (1534-1855) and Myo5B-GTD (1526-1848) proteins were expressed in the *E. coli* BL21(DE3) cells harboring the pET-30-based vectors in LB medium with kanamycin (1 liter each) by induction with 0.1 mM IPTG (16°C, 16 hours). *E. coli* cells were resuspended in 40 ml each of RB150 containing MgCl_2_ (5 mM), DTT (1 mM), PMSF (1 mM), and pepstatin A (1.0 μg/ml), followed by freeze-thaw treatment, sonication, and ultracentrifugation as above. The supernatants were mixed with Complete His-Tag Purification Resin beads (50% slurry, 4 ml each; Roche, Basel, Switzerland) and were incubated at 4°C for 2 hours with gentle agitation to isolate His6-tagged Myo5A-GTD and Myo5B-GTD proteins. The protein-bound beads were washed four times in RB150 containing 5 mM MgCl_2_, 1 mM DTT, and 20 mM imidazole (8 ml each), resuspended in the same buffer (2 ml each) containing HRV 3C protease (15 units/ml; Novagen), and incubated with gentle agitation (4°C, 16 hours). After centrifugation of the bead suspensions (15,300 g, 10 min, 4°C), purified Myo5A-GTD and Myo5B-GTD proteins, which had been cleaved off from the beads, were harvested from the supernatants and dialyzed against RB150 containing 5 mM MgCl_2_ and 1 mM DTT.

Protein concentrations of the purified Rab, HRas, and Myo5-GTD proteins were determined using the Protein Assay CBB Solution (Nacalai Tesque) and BSA as a standard protein. These purified recombinant proteins were boiled (100°C, 5 min) in 1.6% SDS and then analyzed by SDS-PAGE and Coomassie Blue staining.

### GTPase activity assay

GTP-hydrolysis activities of human Rab-family and HRas GTPases were assayed by quantitating released free phosphate molecules during the hydrolytic reactions, using the Malachite Green-based reagent Biomol Green (Enzo Life Sciences, Farmingdale, NY) as described (18, 52), with modifications. Purified recombinant Rab-family and HRas proteins (2 μM final) were incubated at 30°C for 1 hour in RB150 containing 6 mM MgCl_2_, 1 mM DTT, and 1 mM GTP or GTPγS. After a 1-hour incubation, the reaction mixtures (50 μl each) were supplemented with the Biomol Green reagent (50 μl each), further incubated at 30°C for 30 min, and measured for absorbance at 620 nm in a clear 96-well microplate (Falcon no. 351172; Corning, Corning, NY) using the SpectraMax Paradigm plate reader with the ABS-MONO cartridge (Molecular Devices, Sunnyvale, CA). All of the data obtained were corrected by subtracting the absorbance values of the control reactions without any Rab and HRas GTPases. To calculate the concentrations of released phosphate molecules in the reactions, phosphate standard samples (2.5 ‒ 160 μM final; Enzo Life Sciences) were also incubated and assayed using the same protocol. Means and standard deviations of the specific GTPase activities for purified Rab-family and HRas proteins (μMphosphate/min/μM protein) were determined from three independent experiments.

### Liposome preparation

All of the non-fluorescent lipids were purchased from Avanti Polar Lipids (Alabaster, AL). Fluorescent lipids, Rh-PE and FL-PE, were obtained from Molecular Probes (Eugene, OR). Lipid mixes for the Rh-labeled liposomes used in liposome turbidity assays, fluorescence microscopy, and liposome co-sedimentation assays contained 1-palmitoyl-2-oleoyl-PC (POPC) [42% (mol/mol)], POPE (16.5%), soy PI (10%), POPS (5%), cholesterol (20%), DOGS-NTA (6%), and Rh-PE (0.5%). For the biotin/Rh-labeled liposomes and FL-labeled liposomes used in streptavidin bead-based liposome tethering assays and fluorescence microscopy, lipid mixes contained POPC (41%), POPE (14.5% and 16.5% for the biotin/Rh-labeled and FL-labeled liposomes, respectively), soy PI (10%), POPS (5%), cholesterol (20%), DOGS-NTA (6%), biotin-PE (2% for the biotin/Rh-labeled liposomes), and fluorescent lipids (1.5% of Rh-PE and FL-PE for the biotin/Rh-labeled and FL-labeled liposomes, respectively). Dried lipid films harboring these physiological mimic lipid compositions (18, 28) were completely resuspended in RB150 containing 5 mM MgCl_2_ and 1 mM DTT by vortexing, yielding 8 mM total lipids in final, then incubated at 37°C for 1 hour with shaking, freeze-thawed in liquid N_2_ and a water bath at 30°C, and extruded 21 times through polycarbonate filters (pore diameter, 400 or 1000 nm; Avanti Polar Lipids) in a mini-extruder (Avanti Polar Lipids) at 40°C. Lipid concentrations of the extruded liposomes were determined from the fluorescence of Rh-PE (λex = 550 nm, λem = 590 nm) and FL-PE (λex = 486 nm, λem = 529 nm) using the SpectraMax Paradigm plate reader with the TUNE cartridge (Molecular Devices) and a black 384-well plate (Corning no. 3676; Corning). Liposome solutions were diluted with RB150 containing 5 mM MgCl2 and 1 mM DTT to 5 mM total lipids in final and stored at 4°C.

### Liposome turbidity assay

Membrane tethering of Rab GTPase-anchored liposomes was monitored by turbidity changes of liposome solutions as described (18, 25–27), with modifications. After preincubating liposome suspensions and Rab protein solutions separately at 30°C for 10 min, liposomes (400 nm diameter; 0.5 mM total lipids in final) were mixed with Rab proteins (0.5 – 4.0 μM in final) in RB150 containing 5 mM MgCl_2_ and 1 mM DTT (total 150 μl for each), immediately followed by measuring the absorbance changes of the liposome suspensions at 400 nm in a DU720 spectrophotometer (Beckman Coulter) at room temperature for 300 sec. For the assays with class V myosins, Rab proteins (1 μM final) were preincubated at 30°C for 30 min in the presence of Myo5A-GTD or Myo5B-GTD (0.5 – 2.0 μM final) before mixing with liposomes. All liposome turbidity data were obtained from one experiment and were typical of those from more than three independent experiments.

### Liposome co-sedimentation assay

Rh-labeled liposome suspensions (400 nm diameter; 1 mM total lipids in final) were mixed with purified Rab, HRas, or BSA proteins (2 μM in final) in RB150 containing 5 mM MgCl_2_ and 1 mM DTT (100 μl each) and subsequently incubated at 30°C for 30 min. For the experiments with class V myosins, the reaction mixtures were further supplemented with GTP (1 mM final) and Myo5A-GTD or Myo5B-GTD (2 μM final). After incubation, these reactions were ultracentrifuged at 50,000 rpm for 30 min at 4°C with a TLA100 rotor and an Optima TLX ultracentrifuge (Beckman Coulter). The pellets and supernatants obtained were analyzed by SDS-PAGE and Coomassie Blue staining.

### Fluorescence microscopy

After separately preincubating liposome suspensions and Rab protein solutions at 30°C for 10 or 30 min, liposomes (1000 nm diameter; 0.5 or 0.8 mM total lipids in final) and Rab proteins (3.0 or 4.0 μM final) were mixed in RB150 containing 5 mM MgCl_2_ and 1 mM DTT (total 50 μl for each), further incubated at 30°C for 30 min or 1 hour, transferred to ice, and subjected to fluorescence microscopy. For the reactions with class V myosins, Rab11a-His12 (3 μM final) was preincubated in the presence of Myo5A-GTD or Myo5B-GTD (3 μM final) at 30°C for 30 min before incubating with liposomes. A drop of the incubated reactions (5 μl each) was placed on a microscope slide (S2111; Matsunami Glass, Kishiwada, Japan) and covered with an 18-mm coverslip (Matsunami Glass). Fluorescence microscopy was performed using a BZ-9000 fluorescence microscope (Keyence, Osaka, Japan) equipped with Plan Apo 40X NA 0.95 and Plan Apo VC 100X NA 1.40 oil objective lenses (Nikon, Tokyo, Japan) and TRITC and GFP-BP filters (Keyence). Digital images obtained were processed using the BZ-II viewer application (Keyence) and the ImageJ2 software (National Institutes of Health, Bethesda, MD). Particle sizes of Rab-mediated liposome clusters were measured using ImageJ2 after setting the lower intensity threshold level to 100 and the upper intensity threshold level to 255 (19, 53).

### Streptavidin bead-based liposome tethering assay

Liposome tethering assay using a streptavidin-coated bead was performed as described previously (18, 52), with modifications. The biotin/Rh-labeled liposomes (400 nm diameter; 0.08 mM lipids final) and FL-labeled liposomes (400 nm diameter; 0.2 mM lipids final) were incubated with streptavidin-coated magnetic beads (Dynabeads M-280 Streptavidin; Invitrogen) at 30°C for 2 hours, supplemented with the protein solutions containing Rab11a-His12 and Myo5A-GTD or Myo5B-GTD (3 μM each in final), which had been preincubated at 30°C for 30 min, and further incubated at 30°C for 10 min with gentle agitation. The liposome-bound streptavidin beads were washed twice with RB150 containing MgCl_2_ (5 mM) and DTT (1 mM), resuspended in RB150 containing 100 mM β-OG by vortexing to completely solubilize the liposomes on beads, and centrifuged. To quantify the FL-labeled liposomes co-isolated with beads, FL fluorescence (λ excitation = 495 nm, λ emission = 520 nm, emission cut-off = 515 nm) in the supernatants obtained was measured using the SpectraMAX Gemini XPS plate reader (Molecular Devices). Means and standard deviations of the FL fluorescence signals were determined from three independent experiments.

## Acknowledgements

We thank Dr. Naoki Tamura (Institute for Protein Research, Osaka University; now Fukushima Medical University School of Medicine) for valuable suggestions on this project and for substantial contributions to preparing the expression vectors for human Rab GTPases. We are grateful to Drs. Junichi Takagi and Yukiko Matsunaga (Institute for Protein Research, Osaka University) for access to fluorescence microscopy experiments. This study was in part supported by the Program to Disseminate Tenure Tracking System from the Ministry of Education, Culture, Sports, Science and Technology, Japan (MEXT) and Grants-in-Aid for Scientific Research from MEXT (to JM).

## Author contributions

JM designed the research; JM and MI performed the experiments; JM and MI analyzed the data; JM wrote the manuscript.

## Footnotes

This study was in part supported by the Program to Disseminate Tenure Tracking System from the Ministry of Education, Culture, Sports, Science and Technology, Japan (MEXT) and Grants-in-Aid for Scientific Research from MEXT (to JM).

The abbreviations used are: SNARE, soluble *N*-ethylmaleimide-sensitive factor attachment protein receptor; HVR, hypervariable region; DOGS-NTA, 1,2-dioleoyl-sn-glycero-3-{[N-(5-amino-1-carboxypentyl)iminodiacetic acid]-succinyl}); PC, phosphatidylcholine; PE, phosphatidylethanolamine; PI, phosphatidylinositol; PS, phosphatidylserine; Rh, rhodamine; Myo5, class V myosin; GTD, globular tail domain; polymerase chain reaction, PCR; HVR, human rhinovirus; LB, Lysogeny Broth; FL, fluorescein; PO, 1-palmitoyl-2-oleoyl.

---

**Figure S1.**
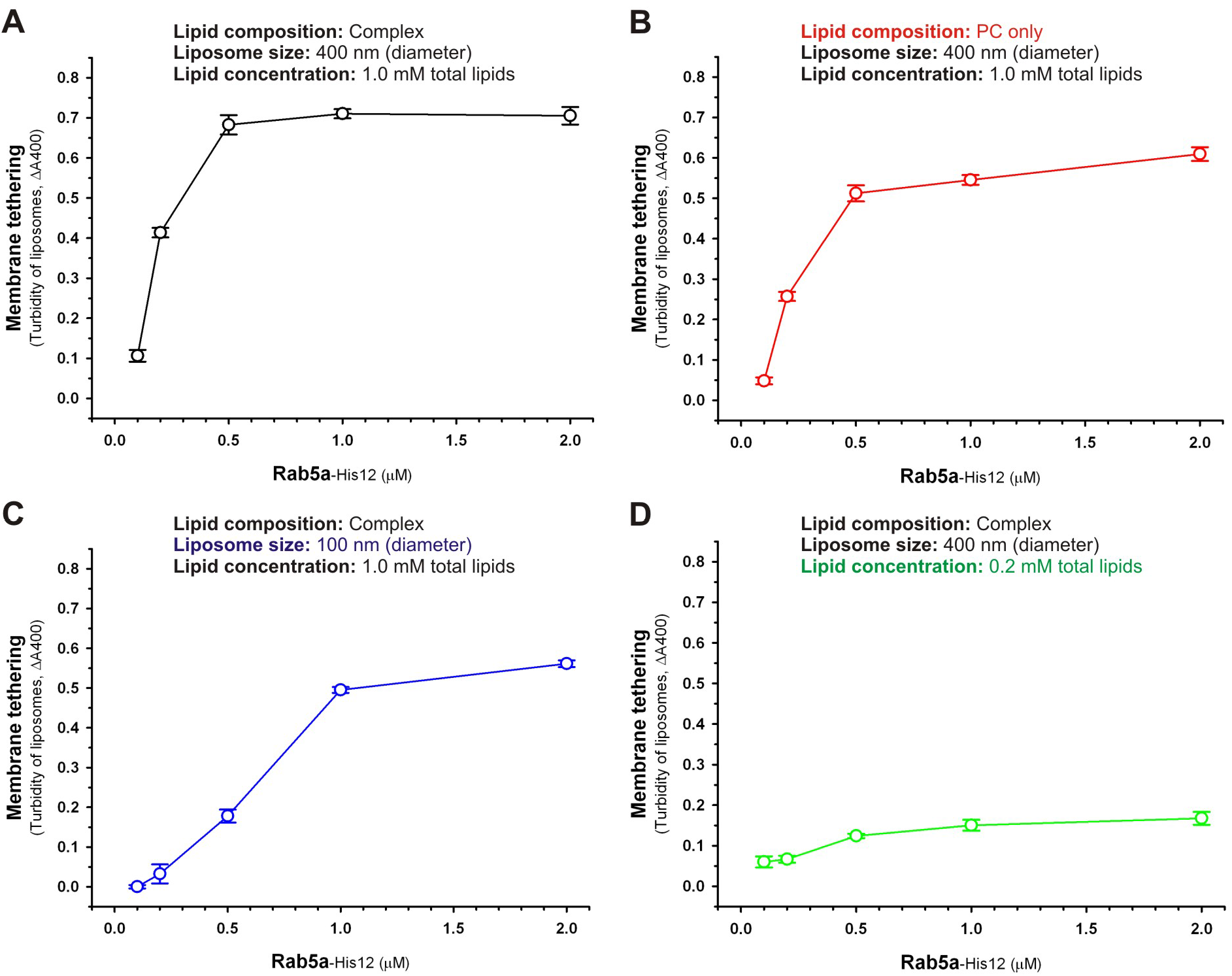
Effects of lipid composition, liposome size, and total lipid concentration on reconstituted liposome tethering mediated by Rab5a-His12. Rab5a-His12 proteins (0.1 – 2.0 (μM final) were mixed with synthetic liposomes in RB150 containing 5 mM MgCl_2_ and 1 mM DTT, and then incubated at 30°C for 30 min. After the 30 min incubation, turbidity changes of the Rab-liposome mixed reactions were measured with the absorbance at 400 nm (ΔA400), using the SpectraMax Paradigm plate reader with the ABS cartridge (Molecular Devices) and a black 384-well plate (Corning no. 3544; Corning). Liposomes used bore the physiological mimic complex lipid composition (Complex) (A, C, D), as in Figure 2, or the PC-only lipid composition [PC only; POPC (92.5%), DOGS-NTA (6%), Rh-PE (1.5%)] (B). Liposome sizes used were 400 nm (A, B, D) or 100 nm (C) in diameter. Total lipid concentrations tested in these turbidity reactions were 1.0 mM (A, B, C) or 0.2 mM (D) in final. Means and standard deviations of the AA400 values were determined from three independent experiments.

**Figure S2.**
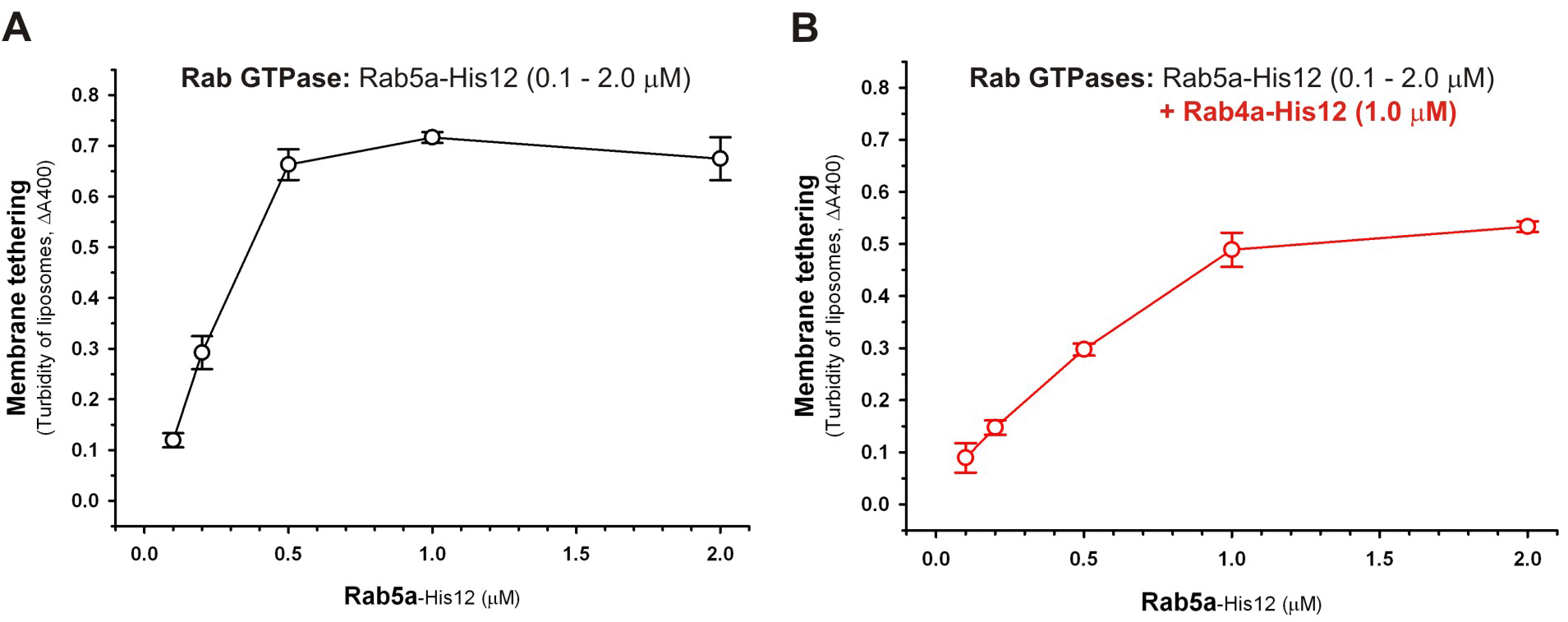
Addition of Rab4a-His12 does not promote liposome tethering mediated by Rab5a-His12. Rab5a-His12 proteins (0.1 – 2.0 (μM final) were mixed with synthetic liposomes and then incubated at 30°C for 30 min as in Figure S1A, but in the absence (A) and presence (B) of Rab4a-His12 (1.0 μM final). After the incubation, turbidity changes of the Rab-liposome mixed reactions were analyzed as in Figure S1.

**Figure S3.**
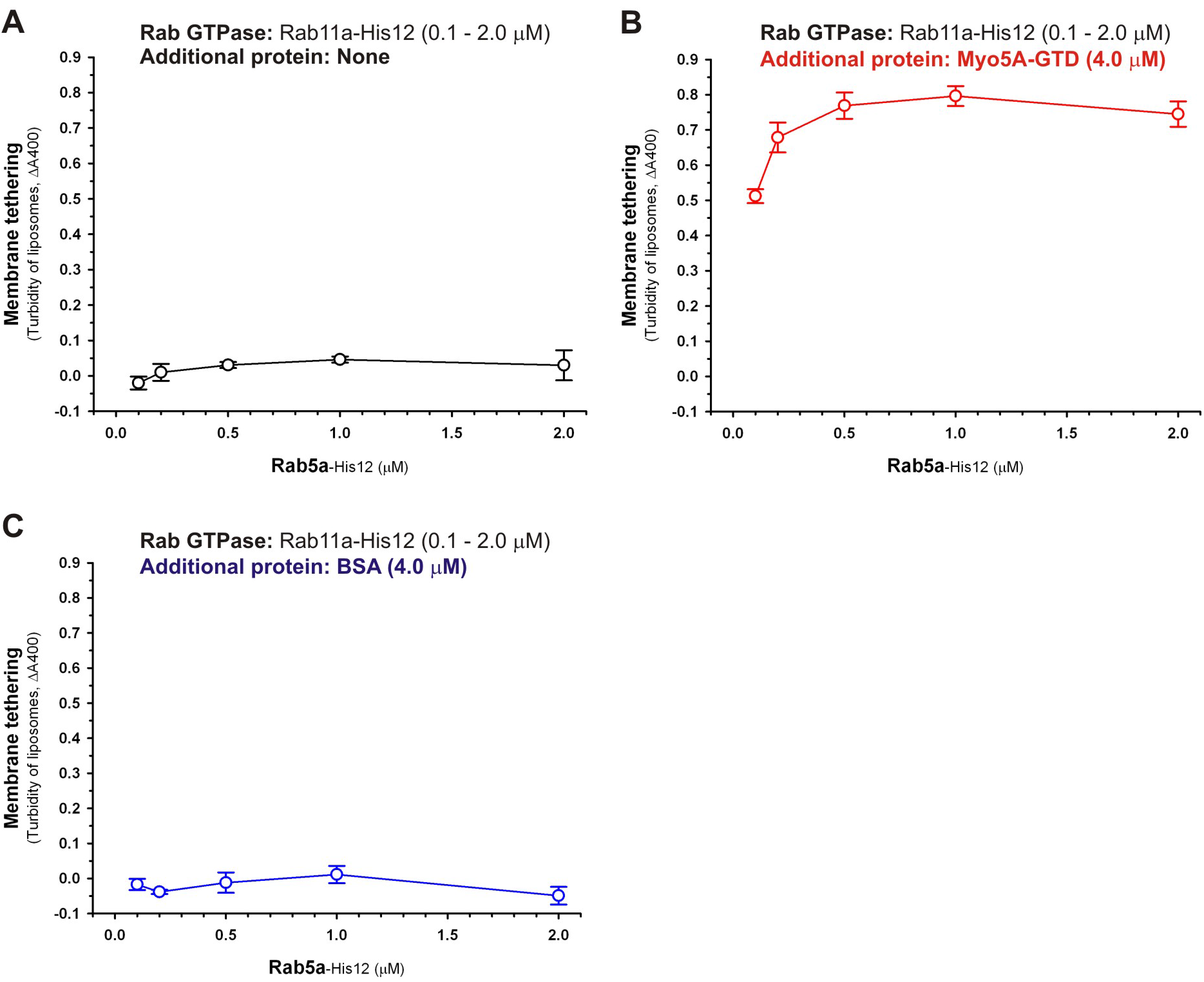
Myo5A-GTD is a specific protein component to promote liposome tethering mediated by the cognate Rab11a-His12. Rab11a-His12 proteins (0.1 – 2.0 μM final) were mixed with synthetic liposomes and then incubated at 30°C for 30 min as in Figure S1A, but in the absence (A) and presence of Myo5A-GTD (4.0 μM final) (B) or BSA (4.0 μM final) (C). After the incubation, turbidity changes of the Rab-liposome mixed reactions were analyzed as in Figure S1.

